# Benchmarking tree and ancestral sequence inference for B cell receptor sequences

**DOI:** 10.1101/307736

**Authors:** Kristian Davidsen, Frederick A. Matsen

## Abstract

B cell receptor sequences evolve during affinity maturation according to a Darwinian process of mutation and selection. Phylogenetic tools are used extensively to reconstruct ancestral sequences and phylogenetic trees from affinity-matured sequences. In addition to using general-purpose phylogenetic methods, researchers have developed new tools to accommodate the special features of B cell sequence evolution. However, the performance of classical phylogenetic techniques in the presence of B cell-specific features is not well understood, nor how much the newer generation of B cell specific tools represent an improvement over classical methods. In this paper we benchmark the performance of classical phylogenetic and new B cell-specific tools when applied to B cell receptor sequences simulated from a forward-time model of B cell receptor affinity maturation towards a mature receptor. We show that the currently used tools vary substantially in terms of tree structure and ancestral sequence inference accuracy. Furthermore, we show that there are still large performance gains to be achieved by modeling the special mutation process of B cell receptors. These conclusions are further strengthened with real data using the rules of isotype switching to count possible violations within each inferred phylogeny.

## Introduction

B cells play a key role in adaptive immunity. After successful VDJ gene recombination of the variable part of the B cell receptor (BCR), and various selection steps, mature B cells are exported from the bone marrow. At this stage the mature B cells have not yet bound antigen and they are therefore referred to as naive. Upon infection some cells from this repertoire of naive BCRs will bind the infectious agent, initializing a cascade of events called affinity maturation leading to pathogen neutralization.

Affinity maturation is a micro-evolutionary process consisting of coupled mutation and selection. This essential process takes place in specialized anatomic compartments called germinal centers (GCs), with the objective of improving antigen binding of the BCR (1). Affinity maturation results in clonal families of thousands of B cells for each of the naive ancestors. Sequences in a family are related to a common naive B cell but with higher affinity BCRs and accumulation of mutations in their sequences.

The study of B cell evolution in the GCs is an important and active field of research including response to infections, mechanisms of vaccines (2) and immunological memory (3). Furthermore, the field has experienced a boost of interest and capability in recent years due to the advancements of high-throughput sequencing of BCR repertoires (Rep-Seq) (4). Rep-Seq now enables sequencing of BCRs on massive scale (millions of cells) and is being increasingly applied in different areas from vaccine studies (5, 6) to antibody engineering (7, 8). Following Rep-Seq, computational methods can be used to group the BCRs into “clonal families”, each consisting of the descendants of a single naive cell (9).

The events of the affinity maturation process can be interrogated by inferring the phylogenies of sequences within each such clonal family, as well as inferring ancestral sequences on the phylogenies. Phylogenetic methods have given great insight into the long and complex development process of broadly-neutralizing antibodies (10, 11). Phylogenetic methods are equally important for shorter-time-scale investigations of affinity maturation, such as of the response to vaccination (12). One may also use trees equipped with ancestral sequences to make statements about the strength of natural selection (13).

Given the importance of these methods to understanding affinity maturation, there has been surprisingly little validation of their performance in the parameter regime relevant to the study of affinity maturation. Although dozens of studies benchmarking phylogenetic methods via simulation in the general phylogenetic case have appeared since (14), methods for BCR sequences deserve special treatment because of special aspects of the evolutionary process of affinity maturation. These include:

1. The somatic hypermutation (SHM) process in affinity maturation is driven by purpose-built molecular machinery (15) that results in a highly context-dependent process with local sequence contexts that either favor (“hotspots”) or disfavor (“coldspots”) mutation (16, 17). The complexity of this process is at odds with both the usual phylogenetic assumption of independent and identical processes between sites and with the assumptions of commonly-used sequence simulators (18, 19) used for benchmarking.
2. Sampling and sequencing, especially for direct sequencing of GCs (20), is dense compared to divergence between sequences. Because the resulting sequences will have limited divergence between them, it raises the possibility that simpler methods with fewer free parameters such as parsimony would be an appropriate choice (21). Also, because of the resulting distribution of short branch lengths, zero-length branches and multifurcations representing simultaneous divergence are common. When these zero-length branches lead to a leaf, they represent a “sampled ancestor” - a sequence with an identical genotype to an ancestral cell. Because of these differences, previous conclusions about performance of phylogenetic estimators in the classical regime of millions of years of divergence need not hold here.
3. Rep-Seq typically sequences the coding sequence of antibodies, which are under very strong selective constraint in GCs. This contrasts strongly with the neutral evolution assumptions of most phylogenetic algorithms, as well as the neutral assumptions of the most common software used for phylogenetics benchmarks (18, 19).
4. In contrast to typical phylogenetic problems where the root sequence is unknown, one has significant information about the root sequence for BCR sequences. Even our current imperfect knowledge of germline genes greatly constrains the space of possible ancestral sequences compared to the typical phylogenetic case where the ancestor is completely unknown. Evolution of BCR sequences happens in a directed fashion from this ancestral sequence.

For these reasons, we believe that BCR-specific validation of phylogenetic tools is an essential prerequisite to their use.

Practitioners frequently use standard phylogenetic tools for BCR sequences. Many studies performing phylogenetic reconstruction on BCR sequences have used the PHYLIP package (22) such as the maximum likelihood (ML) tool dnaml (11, 23–25) or the maximum parsimony (MP) implementation dnapars (26–28). For general phylogenetics use, PHYLIP’s dnaml is now less frequently used compared to faster or more feature-rich programs such as RAxML (29), PhyML (30), FastTree2 (31), and the most recent popular ML program, IQ-TREE (32). However, not all of these programs return ancestral sequence estimates so are less interesting for antibody researchers.

Four tools have been developed specifically for inferring BCR phylogenies: IgTree (33), ARPP (34), IgPhyML (35), and GCtree (36). IgTree aims to find the minimal sequence of events that could have led to the observed sequences (i.e. a maximum parsimony criterion), allowing a known root and sampled ancestors. ARPP is an implementation of a BCR specific ML model to infer ancestral sequences on trees produced by PHYLIP’s dnaml. Both IgTree and ARPP have limited availability: IgTree is not available for download at all, while ARPP is only available for Windows. ARPP cannot be run from a script, thus we could not include it in this large-scale benchmark. IgPhyML adapts the Goldman-Yang (GY94) codon substitution model (37) by adding parameters to model the motif dependent mutation rate. However, to achieve a tractable likelihood the motif contribution is marginalized across codons to achieve a independent-across-codon likelihood function that works well with the usual ML setup. Ig-PhyML is built on codonPhyML (38) which is used for tree sampling and likelihood calculations, ancestral sequence reconstruction can be done in a post processing step using an auxiliary script (provided in the supplement of (35)). GCtree ranks equally parsimonious trees found by PHYLIP’s dnapars according to a likelihood function derived from a Galton-Watson branching process (39). In this branching process, the cellular abundance of a given genotype is used and therefore single cell data is a necessary requirement for optimal ranking with GCtree. Both IgPhyML and GCtree are freely available through GitHub. Additionally, we have implemented an alternative method, called SAMM v0.2, for ranking equally parsimonious trees based on the sum of log likelihoods of the observed mutations between nodes on a tree given a substitution model based on SHM motifs. This ranking is implemented using the SAMM package (40).

To benchmark phylogenetic methods for BCRs, we desired a simulator for full-length BCR sequences that modeled context-sensitive mutation, natural selection on amino acids, and had publicly available source code. Many interesting simulators have different goals. Detailed mechanistic models have been proposed to model all cells and all interactions in a GC using first principles from biophysics (41–43). Others have suggested probabilistic frameworks modeling summary statistics of SHM (44, 45) and, as a middle ground between ultra fine grained models and plain summary statistics, models attempting to explain population level trends using systems of differential equations have been suggested (46). Even simulators that use a notion of sequence don’t necessarily use nucleotides or model mutation in an accurate way. For example, (41) uses a reduced-size alphabet to obtain an appropriately rugged fitness landscape, while (47) use uniform per-site nucleotide mutation in the complementarity determining region and selection based on a subset of key residues.

No existing simulator fit our needs and so we designed a simple model of affinity maturation of BCR sequences in a clonal family. In this model, sequence fitness is solely a function of the amount of antigen bound by the BCR at equilibrium. Antigen binding is calculated using standard binding kinetics applied to a GC with B cells carrying BCRs with different sequences and affinities, competing to bind a limited amount of antigen. Our simple design is motivated by the observation that antigen binding is the main driver and limiting factor of affinity maturation (48). By modularizing the simulation code we have one module preforming mutation and proliferation as a neutral branching process and an optional module to change the birth/death rate through affinity selection.

This simulator has enabled a primary goal of our work: to benchmark methods for ancestral sequence reconstruction. Such methods infer sequences at ancestral nodes of a phylogenetic tree according to some optimality criterion. Ancestral sequence reconstruction is heavily used in BCR sequence analysis, in which it is common to synthesize and test ancestral sequences in order to understand the impact of historical substitutions on binding (49, 50).

A recent and independent effort by (51) did a benchmarking study using simulated BCR sequences without selection and compared phylogenetic method performance, including ML and MP tools. Our study has the following differences with this previous work:

- we simulate sequences under selection using an affinity-based model, which we show makes the inferential problem significantly more difficult,
- we compare accuracy of ancestral sequence inference,
- we include additional software tools, several of which are BCR-specific,
- we provide evidence that our simulations have similar characteristics to real data,
- and we use isotype data as a further non-simulation means of benchmarking methods.

This previous work also worked to understand the results of phylogenetic inference using a “toy” clonal family inference method with necessarily bad performance, whereas here we assume that clonal families have been properly inferred.

In this paper we attempt to answer some of the unresolved questions about BCR phylogenetic inference, including a benchmark of the performance of relevant phylogenetic tools (dnaml, dnapars, IgPhyML, IQ-TREE, GCtree and an undescribed SHM motif based tree ranking method), an investigation of the influence of SHM motifs; and a comparison between simulations with neutral or selection-based evolution (Figure 1). We apply our proposed sequence simulation framework to simulate under different realistic models that include SHM motifs and affinity selection. Finally, we show how the biological mechanism of isotype switching can be used to empirically test phylogenetic inference.

**Figure 1:**
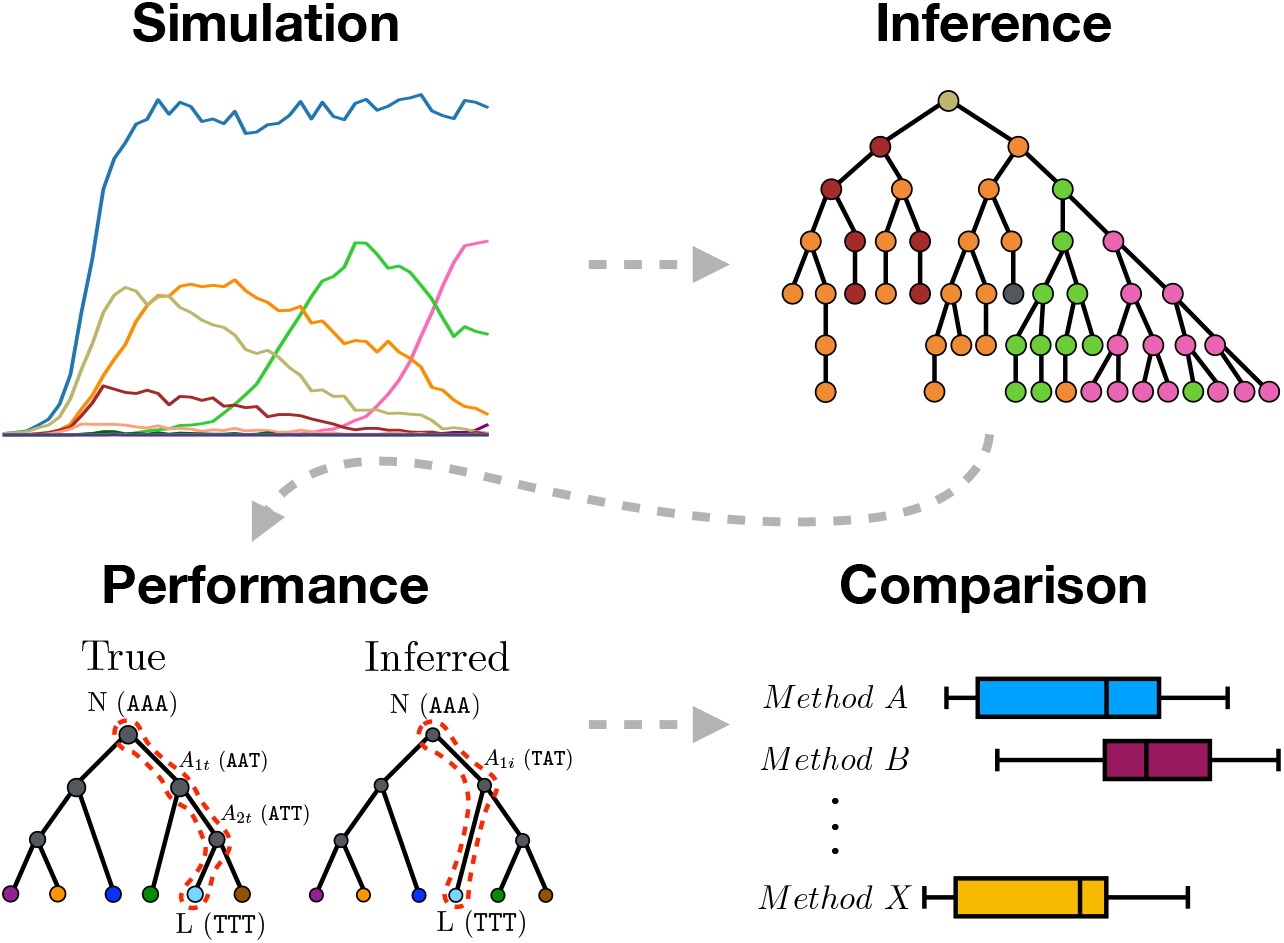
Graphical abstract summarizing the work presented in this paper. We use sequence simulation to establish a ground truth phylogeny from which a sample of sequences is used to infer the phylogeny using different inference methods. The inferred tree is then compared to the simulated true tree to measure inference performance. Lastly, the different inference methods are compared.

All simulation code is open source and can be found on our GitHub repository together with sequence data for the isotype validation (https://github.com/matsengrp/bcr-phylo-benchmark). All simulation data is organized to reproduce figures and is available for download on Zenodo (http://doi.org/10.5281/zenodo.1218140).

## Methods

Although statisticians have made substantial strides in proving identifiability (52, 53) of phylogenetic models and consistency (54) of inferential procedures, proving consistency of phylogenetic methods under context-sensitive BCR evolution models with selection is out of reach because no likelihood function is available. Therefore, we chose the general approach of simulating phylogenies, and benchmark tools based on their inference on samples from these known trees. As ancestral sequence reconstruction is of special interest among the users of BCR phylogenetics (11, 50, 55) we developed a metric to measure ancestral sequence reconstruction performance. In the following subsections we present these simulations and performance metrics, as well as a method to use empirical data to assess performance via the principle of irreversibility of isotype switching.

### Simulation

We devised two simulation strategies for BCR evolution: 1) a neutrally evolving branching process, and 2) a branching process with a birth/death rate controlled by BCR antigen binding. Both simulations start with a single naive sequence as a starting point for the tree simulation; this is evolved a number of generations to a population of BCR sequences from which a sample is drawn and used for inference. To get realistic starting sequences for the simulations we created a set of 288 naive sequences inferred by partis (56) from the healthy donor human single cell dataset in (57) and selected to be of high confidence. When a simulation run is initialized a naive sequence is drawn randomly from this set. Our neutral model is controlled by two parameters which are used to control two Poisson distributions determining the simulation: the progeny distribution (λ) and the mutation generating distribution (λ_mut_). Each evolving sequence has its own λ which expresses the fitness of that sequence in comparison to the other sequences in the population (details below). All sequences have the same mutation probability i.e. λ_mut_ is the same for all sequences and constant throughout the simulation. The simulation starts with a single cell carrying the naive sequence; a draw from Pois(λ) will yield the number of progeny cells in the first generation. If a zero is drawn the cell dies, if one is drawn it propagates without division, if two is drawn it splits into two cells, etc. Next, for each progeny cell a draw from Pois(λ_mut_) will determine how many mutations to introduce into its sequence. Mutations are drawn either from a uniform distribution over both sites and substitutions, or using a context sensitive motif model (e.g. S5F (16)). Multiple mutations are introduced stepwise, one at a time, and if a context sensitive mutation model is chosen the sequence context is updated between each introduced mutation. The simulation process can be terminated in three ways: 1) when all cells have died, 2) at fixed time point *T*, or 3) when a fixed number of cells, *N*, has been reached.

As mentioned above, birth and death rates are controlled through the Poisson rate λ. One can think of this as measuring the level of T helper cell signal, in which lots of signal promotes proliferation while insufficient signal leads to death (1). In our neutral simulations, λ is held constant and is the same for all cells. For simulations with selection we use a very simplistic view of the maturation process, in which selection is purely driven by T helper cell signal which is strong for BCRs binding a lot of antigen and weak for BCRs binding little antigen. To translate this into selection in our simulation framework we devise a simple model to transform a BCR sequence into an affinity value, solve for its antigen binding and then use this to control λ, thus making it sequence dependent. In essence, this “affinity selection” is just a mapping between a BCR sequence and a λ; this enables us to use the same simulation framework for both neutral and affinity simulations.

Here we review the basics of fitness assignment; a detailed description of the model as well as model choices can be found in the supplementary. For any BCR sequence indexed by *i*, its fitness is *λ^(i)^ = Y(x)*, where *Y* is a transformation of some information, *x*, specified in the simulation. For a neutral simulation *Y(x)* is constant and independent of *x*, while for the affinity simulation *Y* is variable with respect to *x*. To model BCR sequence affinity we introduce the concept of a “mature sequence” which is the sequence with the highest attainable fitness in the simulation run. Once the simulation starts the mature sequence acts as an attractor to which evolution tends to converge by rewarding amino acid sequences closer to the attractor with higher λ. The choice of mature sequence is arbitrary so we chose to simulate it by randomly mutating the naive sequence until it accumulates a predefined number of amino acid substitutions. Next, the naive and mature sequence are assigned their own affinity values and the span between these define the affinity gain during affinity maturation. To calculate the affinity of a BCR sequence we calculate its amino acid Hamming distance to the mature sequence and transform this into an affinity value using an appropriate power function calibrated on the naive and mature sequences. We then model the BCR binding kinetics by defining a total GC volume with a constant concentration of antigen and solve for the B cells’ antigen occupancy at equilibrium. Antigen occupancy is mapped to B cell fitness (λ^(*i*)^) using a logistic function returning a value between 0 and 2. These steps describe the general setup of calculating *Y(x)* for the affinity simulation.

Inspection of the simulation runs confirm that affinity simulation recapitulate a number of desired properties (Figure 2: 1) sequence evolution is converging towards the mature sequence, 2) cells are competing for the limited supply of antigen establishing a “carrying capacity”, and 3) favorable mutations are rapidly fixed through selective sweeps (58) analogous to clonal bursts (1, 20). We set the expected number of mutations, introduced into the sequence at each mutation step, to be approximately 0.365. This corresponds to the frequently cited SHM rate at around 10^-3^ (60) given the average length of our naive BCR sequences of 365 nucleotides. We define λ_mut_ = 0.365 as the “normal” mutation rate, but because the estimates of SHM rate vary in the literature we also include half and double of this rate (λ_mut_ ∈ {0.1825,0.365,0.73}) in all our simulations. We observe high correlation between the method performance across all three λ_mut_ (Figure S2 and Figure S3), showing that our conclusions are robust to differences in mutation rate. For neutral simulations the branching parameter (λ) and the population size termination criterion (*N*) are adjusted (λ = 1.5 and *N* = 75) to recapitulate summary statistics of the single cell GC experiment in (20), following a similar procedure as (36). For the affinity simulations the branching parameter is cell-specific and adjusts dynamically, in the range between 0 and 2, according to antigen competition. Each affinity simulation is initialized with 100 mature sequences generated by randomly introducing 5 amino acid substitutions to the naive sequence. Affinity simulations are run with an antigen concentration sufficient to maintain a cell population of approximately 1000 cells, and after 35 generations a random sample of 60 cells is recovered for inference. We also performed intermediate sampling for the affinity simulation: in such cases 30 cells are sampled at generation 15, 30 and 45 and pooled to a total of 90 cells. Neutral simulations were run with 1000 replicates and affinity simulations were run with 500.

**Figure 2:**
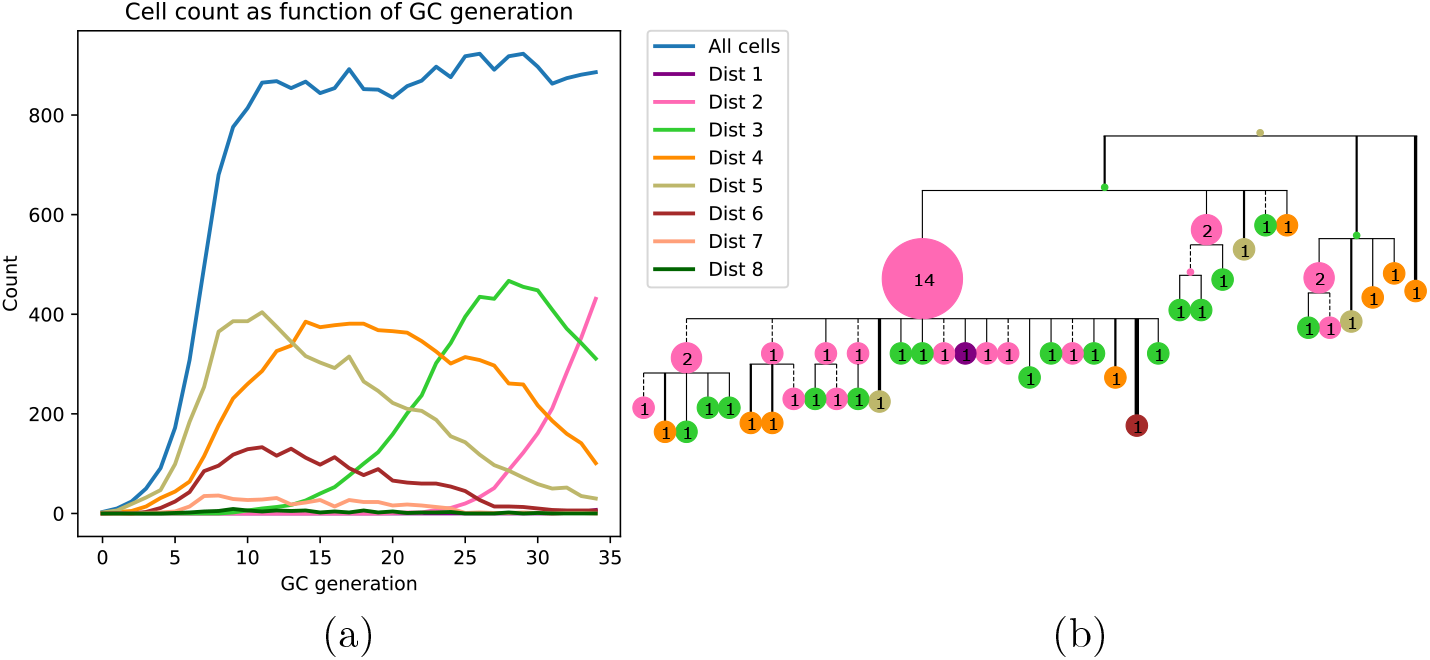
(a) Time series of the distribution of cells at different distances from the mature sequence (Dist 1, 2, …, 8) as the appear in a typical affinity simulation. The simulation is started from a single naive sequence, five amino acid substitutions away from the mature sequence (Dist 5), and simulated sequences converge toward the mature sequence as generations progress. (b) A collapsed tree made from 60 sequences sampled from GC generation 35 of the cell population in (a). Nodes are labeled with numbers indicating the number of collapsed tips (genotype abundance) and node size is proportional to this number. Branch lengths are Hamming distance between nucleotide sequences with dashed lines indicating purely synonymous mutations and solid lines indicating one or more non-synonymous mutations. Branch thickness is proportional to the number of non-synonymous mutations. The tree was rendered with ETE3 (59). Both (a) and (b) are colored according to distance from the mature sequence.

### Inference methods

From each simulation run a subset of sequences was sampled and used for phylogenetic inference along with the correct naive sequence which was used as an outgroup. We tested a number of relevant tools either previously used in the context of BCR phylogenetic inference or with potential use in this field:

- dnaml v3.696: PHYLIP’s implementation of ML using the F84 model (22)
- dnapars v3.696: PHYLIP’s implementation of MP (22)
- GCtree v1.0: Branching process likelihood ranking of MP trees (36)
- SAMM v0.2: Mutation motif based likelihood ranking of MP trees (40)
- IgPhyML v0.99: GY94 codon model with hot/cold spot motif parameters
- (35)
- IQ-TREE v1.6.beta5 (IQT): Fast ML inference with many substitution models (32)

For all methods the naive sequence was used as an outgroup, furthermore, the naive sequence was used to reroot the tree after inference. IQ-TREE was run using either JC, HKY or GTR nucleotide substitution models and using the “ASR” flag, but otherwise with default settings. IgPhyML was run as described in (35) and using the “`-o tlr -s`” flags to optimize both branch lengths and topology under the HLP17 model. dnaml was run using gamma distributed rates, a coefficient of variation of substitution rate among sites of 1.41, four rate categories and otherwise default parameters. dnapars was run using default settings. In the case of dnapars it is common to observe many equally parsimonious trees, and in those cases a random tree was drawn. GCtree was run as described in (36), passing both sequences and their abundances to the program. Both GCtree and SAMM use the equally parsimonious trees generated with dnapars for likelihood ranking, hence in the case when only a single MP tree is found, dnapars, GCtree and SAMM will by definition yield the same result.

The use of all the above methods has been described previously, except SAMM which is part of a statical framework to infer DNA mutation motifs using survival analysis (40). As it is well known that SHM is context sensitive (16, 17, 61) we attempted to use the idea from (36) but ranking equally parsimonious trees according to their SHM motif likelihood rather than a branching process likelihood. Using SAMM we calculate the likelihood of the observed mutations given a tree equipped with ancestral sequences at the internal nodes (in this application from parsimony) and a motif model by using Chib’s method (62) to integrate out event orders on the branches.

We would like to make it very clear that we use the same motif model for both simulating mutations and calculating SAMM likelihoods. This gives SAMM an unfair advantage, however, the selection process is not modeled as part of the motif model. We are not formally proposing SAMM ranking as a competing inference method, but rather as a yardstick with which to measure how much improvement would be possible taking a fully context-sensitive mutation process into account. On the other hand, SAMM has no inherent advantage on the isotype scoring experiment, and it is limited to the MP trees.

### Genotype collapsing

Due to our focus on ancestral sequence inference we have adopted the use of genotype collapsed trees from (36) throughout this work. Briefly, a genotype collapsed tree is made by inferring a phylogenetic tree, inferring ancestral sequences at the internal nodes and recalculating the branch lengths as Hamming distances between the node sequences. In the branch length recalculation step nodes are “collapsed” if their sequences are identical, thereby collapsing tips upwards and adding observations to internal nodes (Figure 2, b). Genotype collapsing deals conveniently with the very short branch lengths, typically observed in binary trees for BCR sequences, since these most often collapse into a single node.

### Tree metrics

We scored trees both in terms of tree structure and in terms of ancestral sequence inference. For tree structure, we used the commonly used Robinson-Foulds (RF) distance (63), which is half the size of the symmetric difference between the sets of bipartitions obtained by cutting each edge. We define bipartitions using both tips and sampled internal nodes, as opposed to standard RF using only tips. Because we perform RF on genotype-collapsed trees, this measure in fact combines accuracy estimation of ancestral sequences and tree topology.

We also used several means to more directly compare ancestral sequence reconstructions: the “most recent common ancestor” (MRCA) metric, and the “correctness of ancestral reconstruction” (COAR) metric. The MRCA metric compares ancestral sequences on the true vs. the inferred phylogeny in a way that does not depend on agreement between the two topologies. Specifically, the MRCA distance is calculated by iterating through all pairs of leaves. For each such pair there is a well defined MRCA node on the tree. The MRCA metric is the average Hamming distance between the inferred and the true ancestral sequence for these pairs. Using *i* and *j* (*i* ≠ *j*) to iterate over all combinations of pairs of leaves to find their true (*T_i,j_*) and inferred (*I_i,j_*) most recent common ancestor, this can be written as:

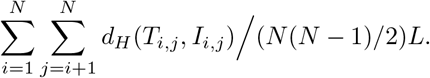

Here *N* is the number of leaves and *L* is the length of the sequence. Thus, MRCA gives an overall view of how ancestral sequence reconstruction is performing.

There is also a special interest in benchmarking tools to reconstruct a lineage of ancestral sequences going from the root (the naive sequence) to a tip of interest (11, 55). Hence, we developed the COAR metric which is measuring the average number of sequence mismatches across all true vs. inferred lineages going from the root to any tip. It is not initially obvious how to compute such a distance if the true and inferred lineage contains a different number of nodes. We solve this problem by finding the node to node comparison that minimizes the distance while maintaining the root-to-tip order. Please see the Supplementary Information for details on COAR metric calculation.

We chose COAR as our principal metric for comparison because it was well correlated with other metrics (see Results) and because it reflects how researchers use ancestral sequence reconstruction of BCRs.

### Isotype scoring

We used sequences with isotype information as another means of characterizing phylogenetic accuracy. The isotype-determining constant region is located downstream of the heavy chain BCR variable region, and isotype changes through a process called class-switch recombination. In mice the isotype constant regions are ordered, from closest to furthest to the J gene: IgM, IgG, IgE, then IgA. Naive BCRs use IgM, but during affinity maturation isotype switching can occur by looping out one or more of the constant regions. For instance if IgM is looped out the resulting BCR is IgG and if IgM, IgG and IgE is looped out the resulting BCR is IgA. Because the isotype is physically removed from the chromosome this process is irreversible, hence a parent cell with an IgA BCR can never give rise to a child cell of IgM isotype.

We use the irreversible nature of isotype switching to measure the performance of tree inference by mapping back isotype labels to the nodes on the inferred tree and counting the number of nodes with an edge to a child that violate the rules of isotype switching. We use the BCR data from (64) which is generated with UMI technology and primers targeting the isotype region on splenocyte whole mRNA from five outbred mice undergoing an immunization campaign. After extensive quality filtering using pRESTO (65) we ran partis (9) to partition sequences into clonal families. These clonal families were filtered based on having minimum 10 and maximum 200 unique sequences and containing at least two different isotypes. Furthermore, we discarded all clonal families where inference exceeded 24 hours of compute time for any single tool on a single core. This left 697 clonal families to do isotype validation.

We defined an isotype mismatch as an observed violation of the isotype switching order (namely the order IgM, IgG, IgE, IgA). That is, an edge connecting a parent and a child node is an isotype mismatch if the isotype order of the parent is farther along the order than its child (Figure S13). To calculate the “isotype score” we iterate over all the tips and use each tip as a starting point to collect the list of isotypes between this tip and the root. This list is made by progressing from a tip to the root and collecting isotypes sequentially, however, unobserved internal nodes will not have an associated isotype and therefore they “reverse inherit” the isotype from their child. Once this list has been filled, each edge is evaluated and if an isotype mismatch is encountered the parent node is marked as a violator. The number of isotype switching violations is found by counting all the violator nodes.

This sum is dependent upon the shape of the inferred tree, potentially leading to a bias associated with each inference tool. To address this, for each inferred tree we created 10,000 samples of trees with the same topology but shuffled labels and from these we calculated a “baseline” isotype score to be expected given this topology. We divided the violation count by the baseline to obtain the final isotype score.

### Boxplot layout

Tool performance is plotted in boxplots. Colored boxes cover from lower to upper quartiles, with the median marked by grey vertical lines and whiskers extending to 1.5 times the interquartile range. Points beyond the range of the whiskers (outliers) are hidden for clarity. Red triangles mark the mean metric value of all simulations, with 1000 replicates for neutral and 500 replicates for affinity simulations, with an overlapping horizontal red line showing the 95% confidence interval of the mean. Confidence intervals were computed using non-parametric bootstrapping, using sampling with replacement to generate 10,000 bootstrap replicates (66). Tools are ordered according to their mean metric values.

## Results

### Metrics are correlated

The RF, MRCA and COAR metrics are highly correlated, with COAR being the most central metric (Figure 3). We checked this for both neutral and affinity simulation and over a range of mutation parameters (Figure S1) and conclude that the high correlation between metrics is robust over many parameter choices. To reduce the number of comparisons we chose COAR as our principal metric because this was the most central metric as well as being interpretable as the expected number of per-site errors per reconstructed lineage. However, all metrics have been run on all simulations (see supplementary figures), except RF distance which does not deal well with reoccurring sequences that appear multiple times in the affinity simulation.

**Figure 3:**
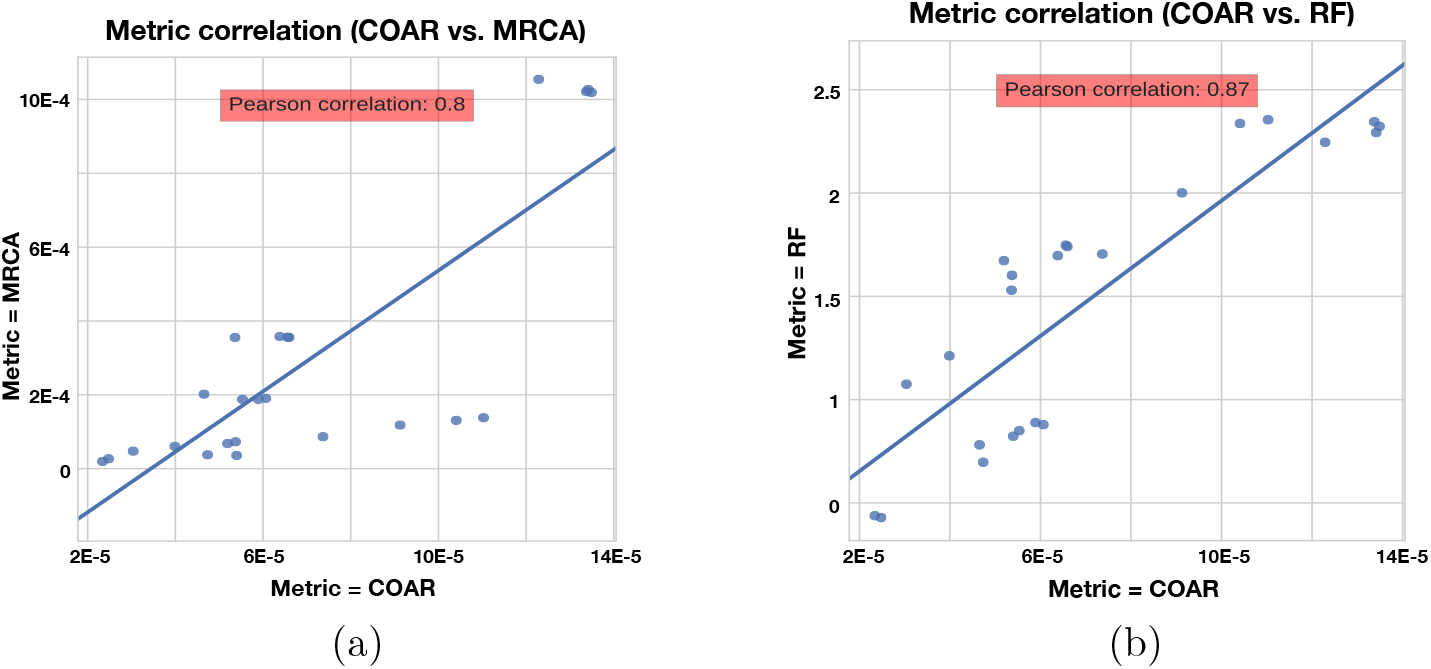
Correlation between metrics for the neutral simulation across the three mutation rates described in Results. Same trend is true for affinity simulation (Figure S1).

### Methods differ in performance consistently across simulations

We observe similar trends across varying simulation methods, performance metrics and mutation rates. A higher mutation burden (λ_mut_) leads to more complex trees resulting in decreased inference performance, and this is true for all methods and performance metrics (Figure S4 to Figure S10). Tools perform better on neutral simulation compared to affinity simulations (Figure 4), which is to be expected due to the added complexity of the affinity simulation. Overall, the distributions of performance metrics are heavy tailed with several outliers far outside of the interquartile range. We have chosen to hide such outliers for the interpretability of our boxplots but their impact can be observed in the means (red triangles) and their confidence intervals.

**Figure 4:**
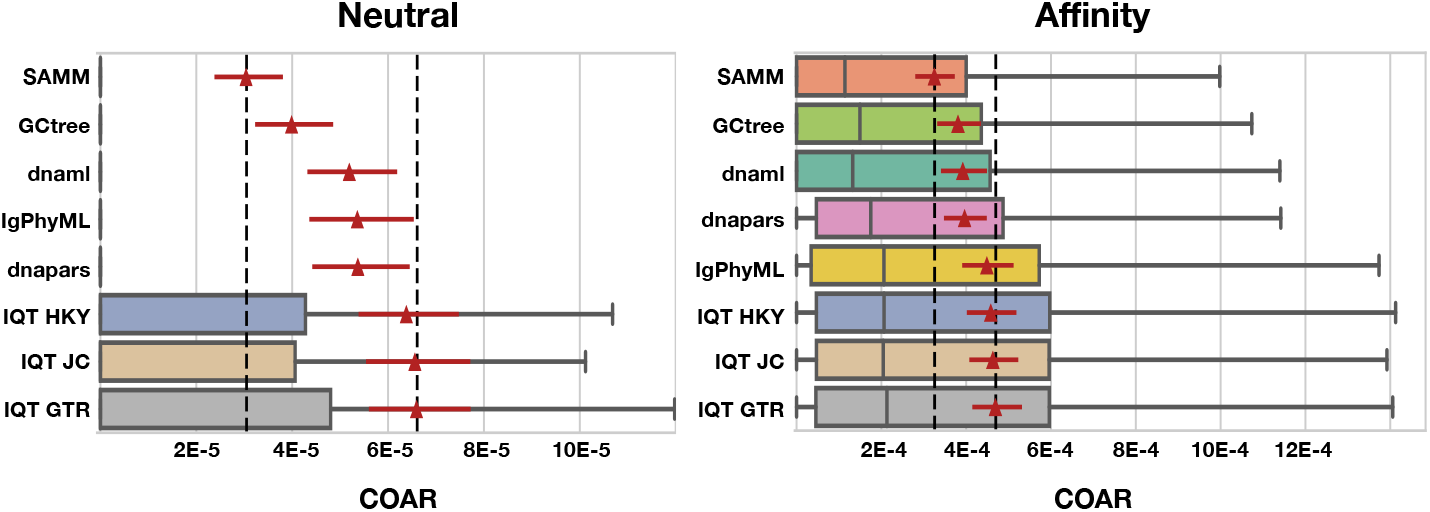
COAR performance for different tools under neutral and affinity simulation using normal SHM rate (λ_mut_ = 0.365) and mutations drawn from the S5F motif model. Colored boxes cover the lower to the upper quartiles, with the median marked by grey vertical lines and whiskers extending to 1.5 times the interquartile range. Points beyond the whiskers (outliers) are hidden for clarity. Red triangles mark the mean COAR value of all simulations (1000 replicates for neutral and 500 replicates for affinity simulations) with the overlying red lines showing the 95% confidence interval found by bootstrapping with 10,000 replicates. Black dashed lines mark highest and lowest mean COAR values. Tools are ordered according to their mean COAR value.

We find that SAMM and GCtree, which rank equally-parsimonious trees, perform better than a uniformly-selected equally parsimonious tree from dna-pars. For all 15 tests across mutation rates, performance metrics and simulation methods SAMM is better than dnapars while GCtree is better than dnapars 13/15 times (Figure S4 to Figure S10). SAMM is the best ranked tool 12/15 times and often with a substantial margin to the second best. Thus the equally-parsimonious tree set contains better and worse trees, and the likelihood ranking of these is effective at distinguishing between them. However, given that SAMM were using the S5F model for likelihood calculations on simulated mutations also drawn from an S5F motif model, it should be not surprise to see that SAMM consistently outperforms all other tools.

Because SAMM is constrained by dnapars and the criterion of only ranking equally parsimonious trees, we consider the performance of SAMM compared to other tools as a conservative estimate of the potential improvement available when correctly modeling SHM motif bias. As a control, we note that when mutations are drawn from a uniform distribution over sites and substitutions, SAMM is not any better than dnapars (Figure S11 and Figure S12) showing that SAMM’s performance can be ascribed to the mutational context bias. Thus, we can use the performance difference between SAMM and dnapars to measure how much inference performance can improve by incorporating SHM motif bias.

Simulated datasets include information on sequence abundance, which enables good performance of the GCtree method. Normally, phylogenetic trees are made from a set of unique sequences while the cellular abundance of each sequence, referred to as genotype abundance, is discarded. GCtree, on the other hand, utilizes this genotype abundance information by ranking equally parsimonious trees via a likelihood using abundances. Our results show that GCtree is the second best performing tool, and consistently better than picking a random equally parsimonious tree, indicating that the integration of genotype abundance information does improve tree inference. Here GCtree is given the correct abundances, giving an upper bound on the performance gain obtainable by incorporating abundance information. In a situation with real data GCtree would rely on single cell data to gain estimates of genotype abundances; while single cell data is becoming more widespread (57, 67–69) the majority of Rep-Seq studies are still based on bulk RNA sequencing resulting in unknown genotype abundances.

Performing third best after SAMM and GCtree comes dnaml and dnapars, both with similar performance, after that IgPhyML and lastly the three mutation models implemented in IQ-TREE which are all performing very similarly (Figure 4). dnapars performs slightly better than dnaml in neutral simulations while the opposite is true in affinity simulations. Practically, the difference between the two programs is so small that we suggest users to choose whichever program they find to be fastest or most convenient to use for their application.

Surprisingly, on simulated sequences IgPhyML performs consistently worse than the simpler dnaml or dnapars alternatives. Although, it is clear from the SAMM results that SHM motifs are present and provide useful information for inference, it does not seem to improve IgPhyML performance beyond SHM naive methods such as MP. IgPhyML’s model was preferred (by likelihood ratio test) in the examples provided in the paper introducing it, which were large trees of long-term broadly-neutralizing anti-HIV antibodies (35). We suspect that IgPhyML’s model is too rich for the less complex data provided here.

All three IQ-TREE methods, using different mutation models, perform consistently worse than any other tool tested in this study. We find it surprising that IQ-TREE using the HKY model is so far off dnaml using F84 despite the high similarity between the two substitution models. We therefore conclude that implementation differences e.g. tree space search, convergence criteria etc. must be the reason for this discrepancy, which is in concordance with our observation that IQ-TREE is much faster than dnaml.

### Isotype data confirms that raw parsimony is worse than likelihood models

The results of our investigation using isotype were somewhat inconclusive. This measure had an extraordinarily large variance observed in both the confidence intervals and the changed rankings upon rerunning the analysis (Figure S14). Although SAMM did perform best among all tools when using a custom motif model fitted on the whole isotype dataset, the difference to other tools was small relative to the variance. Despite this, SAMM is significantly better than dnapars (Figure 5), again confirming the notion that the SHM mutation process is important and contains residual information not captured by the parsimony objective. Notably, the parsimony ranking of GCtree is also significantly better than dnapars (Figure S14) despite the fact that this dataset did not contain genotype abundance information. This indicates that the branching process prior used by GCtree can also yield useful results using the tree topology alone. Testing the full potential of GCtree would require a single cell dataset and this may also result in even better performance.

**Figure 5:**
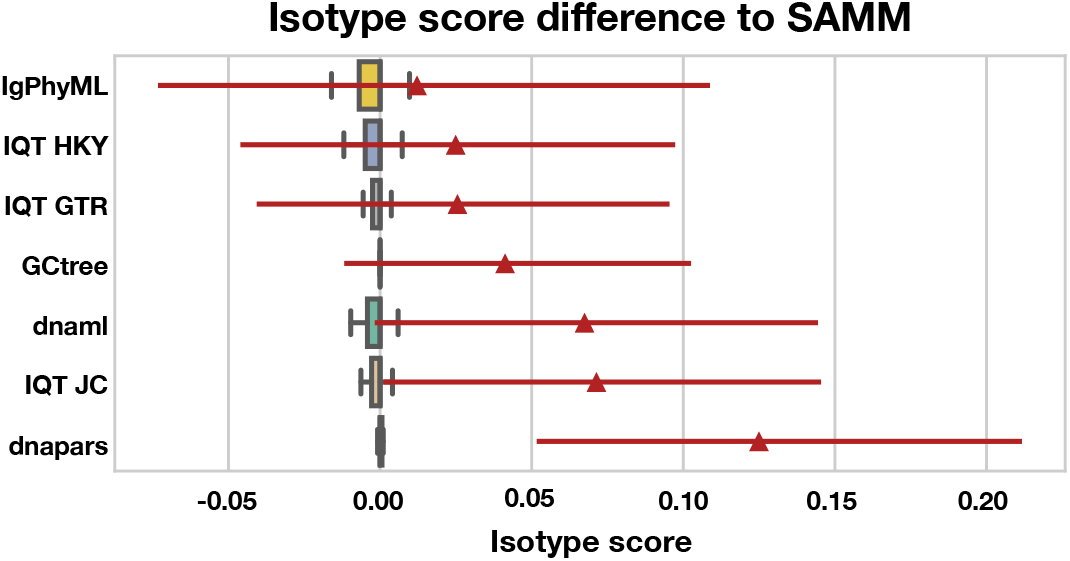
Isotype score difference to SAMM for 697 clonal families with isotype information. Positive differences mean higher (worse) isotype score than SAMM and vice versa for negative values. The 95% confidence interval of the mean difference to dnapars is positive and non-zero showing that SAMM is significantly better than dnapars.

## Discussion

In this work we have benchmarked the performance of phylogenetic algorithms for use in B cell sequence analysis, with a special emphasis on ancestral sequence reconstruction. Our sequence simulation deviates from the standard independent-across-nucleotides models, often used in such benchmarking, by both introducing mutations using a realistic SHM motif model and rewarding convergent mutations via an affinity model of the binding equilibrium between BCRs and antigen. To our knowledge this is the first simulation method to model affinity maturation using BCRs represented as DNA sequences such that selection is based on the corresponding amino acid sequences. Inference based on affinity simulated sequences is more challenging, resulting in ~10 fold higher COAR values (Figure 4), underlining the importance of considering selection to get realistic error estimates on BCR phylogenetic reconstruction. Still, the average COAR values for affinity simulation is 0.0003-0.0005 which translates to an expectation of 1-2 total nucleotide errors in a lineage with 5 heavy+light chain BCR sequences reconstructed (~3600 nucleotides). With the added benefit that about 1/3 of these expected mutations will be silent, reconstruction of BCR affinity matured lineages using ancestral sequence reconstruction in this parameter regime appears to be of high fidelity. However, this estimate should be tempered with the fact that the correct naive sequence was provided to the algorithm, and the general fact that complex processes happening in real data can make the problem significantly harder.

Looking at the more subtle differences between tools two observations stand out: first, accounting for SHM motifs is the biggest contributor to accuracy, and second, implementation matters. The performance of SAMM on simulations clearly shows how SHM motifs leave a useful trace that can be integrated into an inference method. One such method is the HLP17 model used by IgPhyML (35), but it may suffer from noisy parameter estimates in cases with relatively few sequences per clonal family. An extension to IgPhyML may alleviate these problems by either fixing the hot/cold spot parameters with a predetermined motif model, or the means of combining information across clonal families. Yet, there are still reasons to attempt other ways of integrating SHM motifs, as well as other affinity maturation specific information like genotype abundances, into inference methods in more principled ways than mean field approximations or likelihood ranking of MP trees. Our benchmark also gives a reminder that implementation matters. Under otherwise similar substitution models two different implementations (dnaml and IQ-TREE) vary substantially and consistently in performance. We do not know what causes these differences, but we speculate that tree space sampling could be a critical point as this appears to be the most important difference between these two implementations, and because IQ-TREE experiences the same pathologies with multiple different substitution models. IQ-TREE’s heuristics were probably tuned with the traditional phylogenetic case (of deeply diverging sequences) in mind, which is different from our use case.

BCR isotype switching is an irreversible event and contains useful information about the phylogenetic relationship among BCR sequences in the same clonal family. We observed that the two MP tree ranking methods (SAMM and GCtree) did significantly decrease the isotype score compared to picking a random equally parsimonious tree, thus confirming our simulations. Despite this it appears to be very difficult to use the isotype score as an empirical performance metric because of its high variance. We believe that this is in part due to sparse sampling of the clonal families (only few tens of sequences out of the thousands evolved in a GC). With better sampling and more clonal families we expect the isotype score to be better resolved, with lower variance, and then it may be a more useful metric for assessing the performance of BCR phylogenetic inference, or simply used as a constraint in the inference model itself (70).

In this work we provided phylogenetic algorithms with the correct naive sequence. The impact of naive sequence uncertainty was in a way benchmarked by (51), in which they used a coarse method for clonal family inference and then asked if phylogenetic methods could later disentangle the families. Both our study and (51) leave open the question of the performance of phylogenetic methods when supplied with a potentially noisy estimate of the naive sequence supplied by current clonal family inference tools. We will perform the appropriate benchmarking as part of our future development of methods to perform phylogenetic reconstruction and naive sequence estimation simultaneously.

In this work we also have not tested the impact of insertion-deletion (indel) mutations, which do happen in BCR phylogenies (61, 71, 72). Current tools leave a lot to be desired for ancestral sequence inference in the presence of indels, as in our experience they “fill in” nucleotides at every site of an ancestral sequence inference, even if a gap is clearly the right choice. In addition, indels are not treated as the informative characters they are in mainstream phylogenetics software; rather, they are treated as missing data. Benchmarking phylogenetic tools would also require benchmarking the alignment step, which has an effect on ancestral sequence reconstruction accuracy (73). Nevertheless, this will be another important focus for future tool development and ancestral sequence reconstruction benchmarking within the field of BCR phylogenetic reconstruction.

## Acknowledgments

We would like to thank Francois Vigneault and Andreas Laustsen for sharing the mouse B cell receptor sequencing dataset from (64), and David A. Shaw for preparing mutability and substitution matrices specific for this dataset using SAMM, and for providing the SAMM-rank code. Our simulation framework relies on code developed by William DeWitt and was greatly improved by comments and suggestions from Amrit Dhar and Vladimir Minin. This research was supported by National Institutes of Health grants R01 GM113246, R01 AI120961, and U19 AI117891. The research of Frederick Matsen was supported in part by a Faculty Scholar grant from the Howard Hughes Medical Institute and the Simons Foundation.

## Supplementary Materials

**Figure S1:**
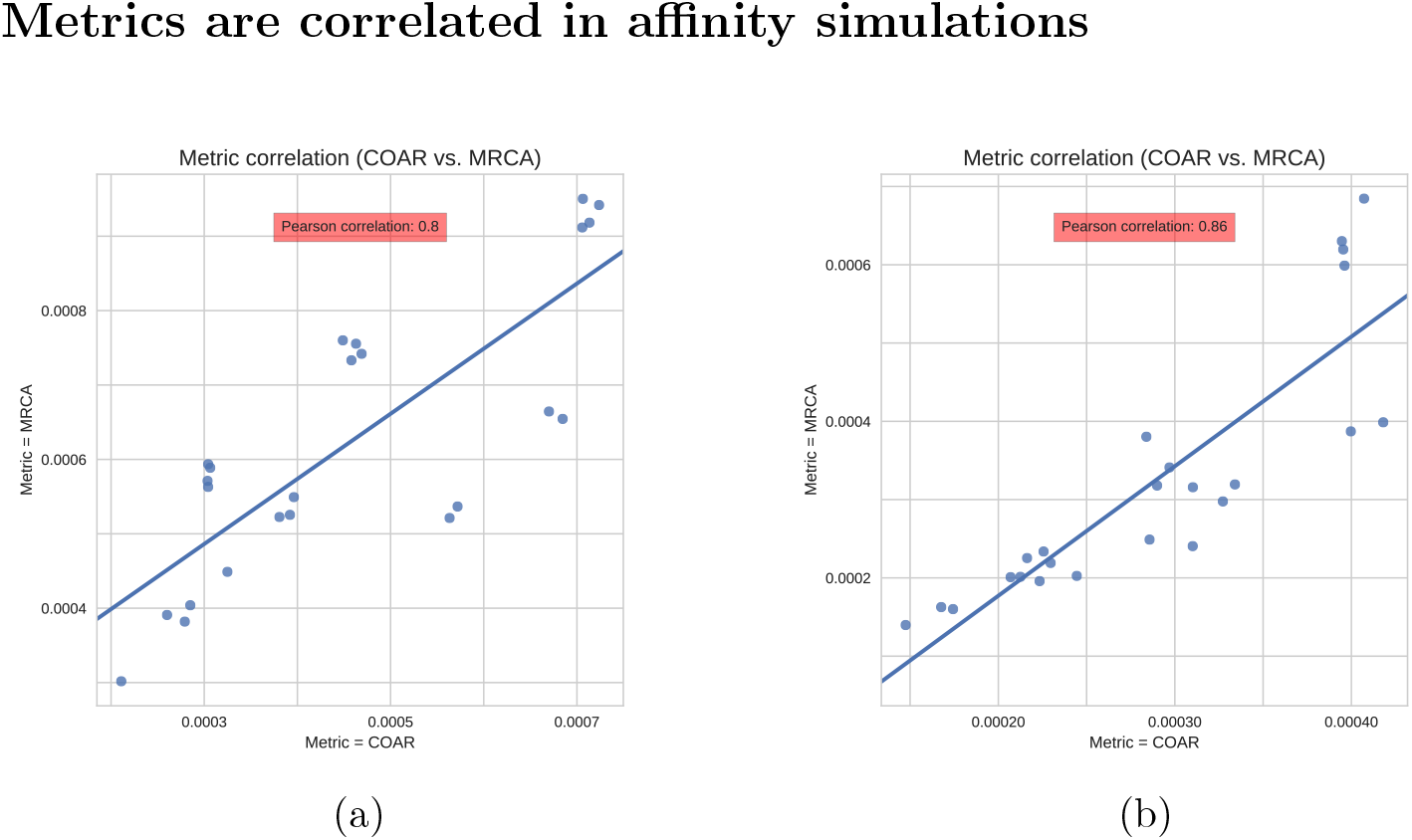
Metric correlations for affinity simulations across three different mutations rates (∀λ_mut_ ∈ {0.1825,0.365,0.73}). a) Single sample. b) Three samples, with two intermediate sampling times.

**Figure S2:**
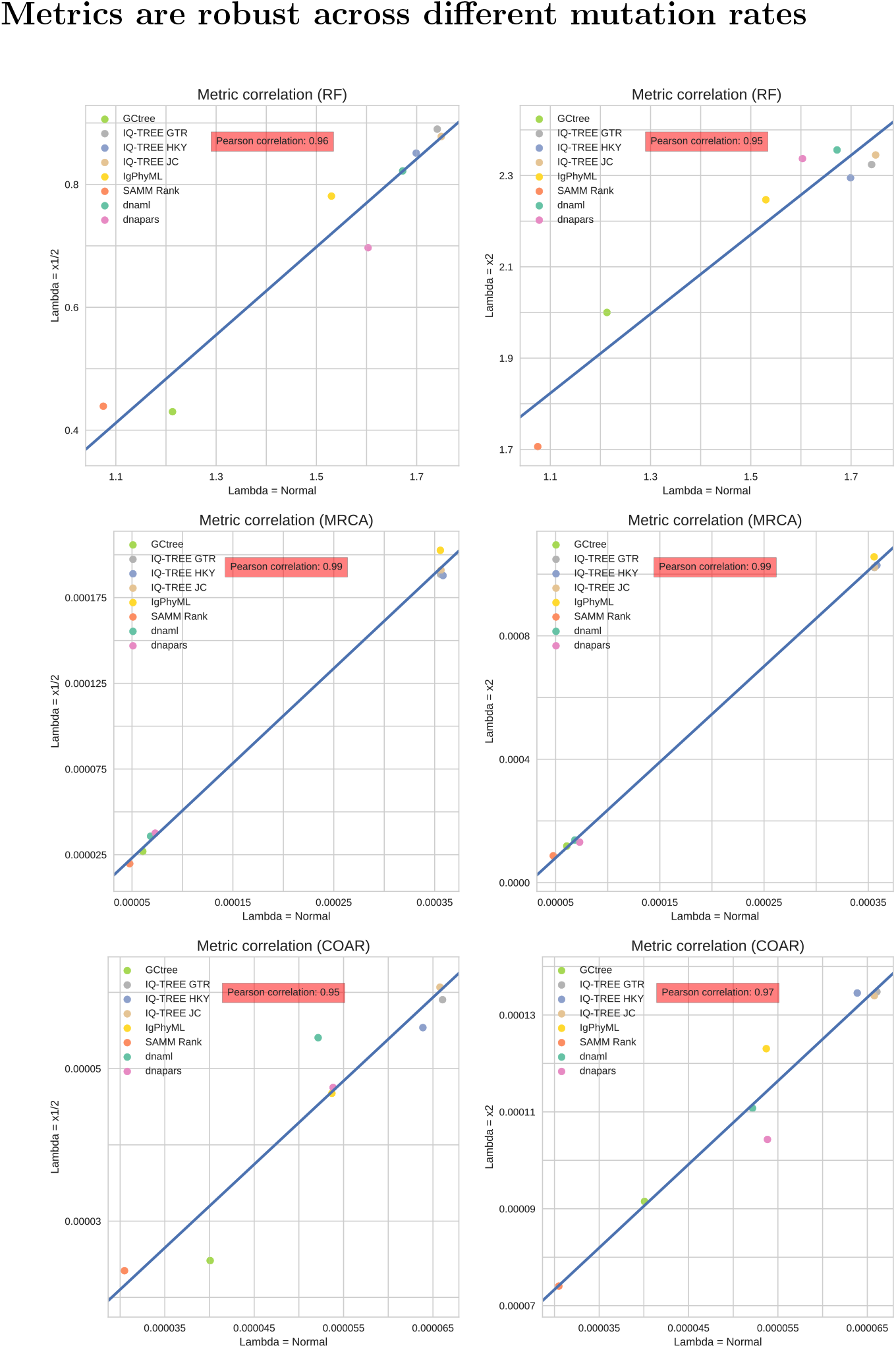
Correlation between the average performance of the methods tested at different mutation rates for neutral simulations over all three performance metrics.

**Figure S3:**
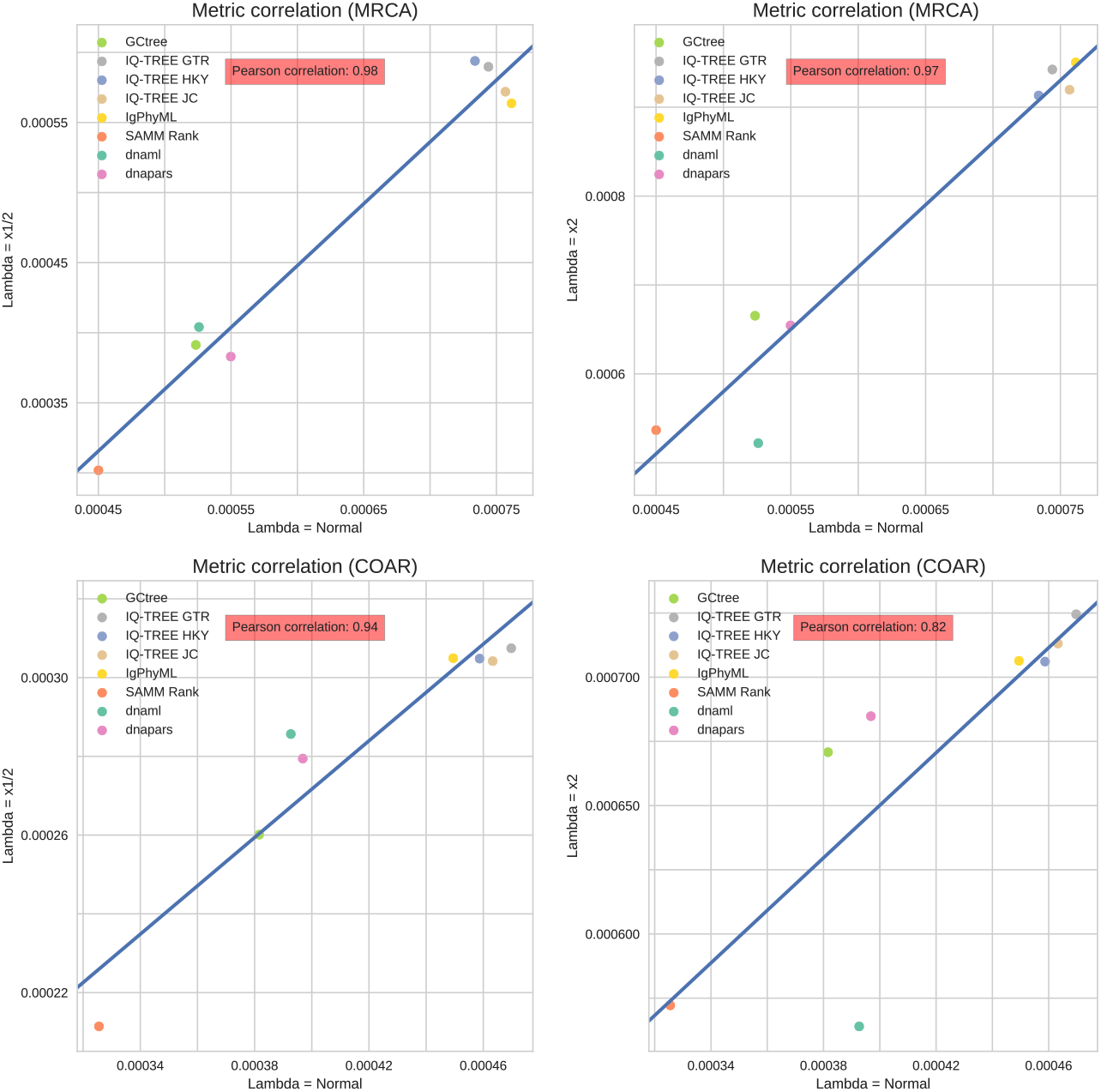
Correlation between the average performance of the methods tested at different mutation rates for affinity simulations over the two performance metrics (RF distance excluded because of recurring sequences in the simulated phylogeny).

**Figure S4:**
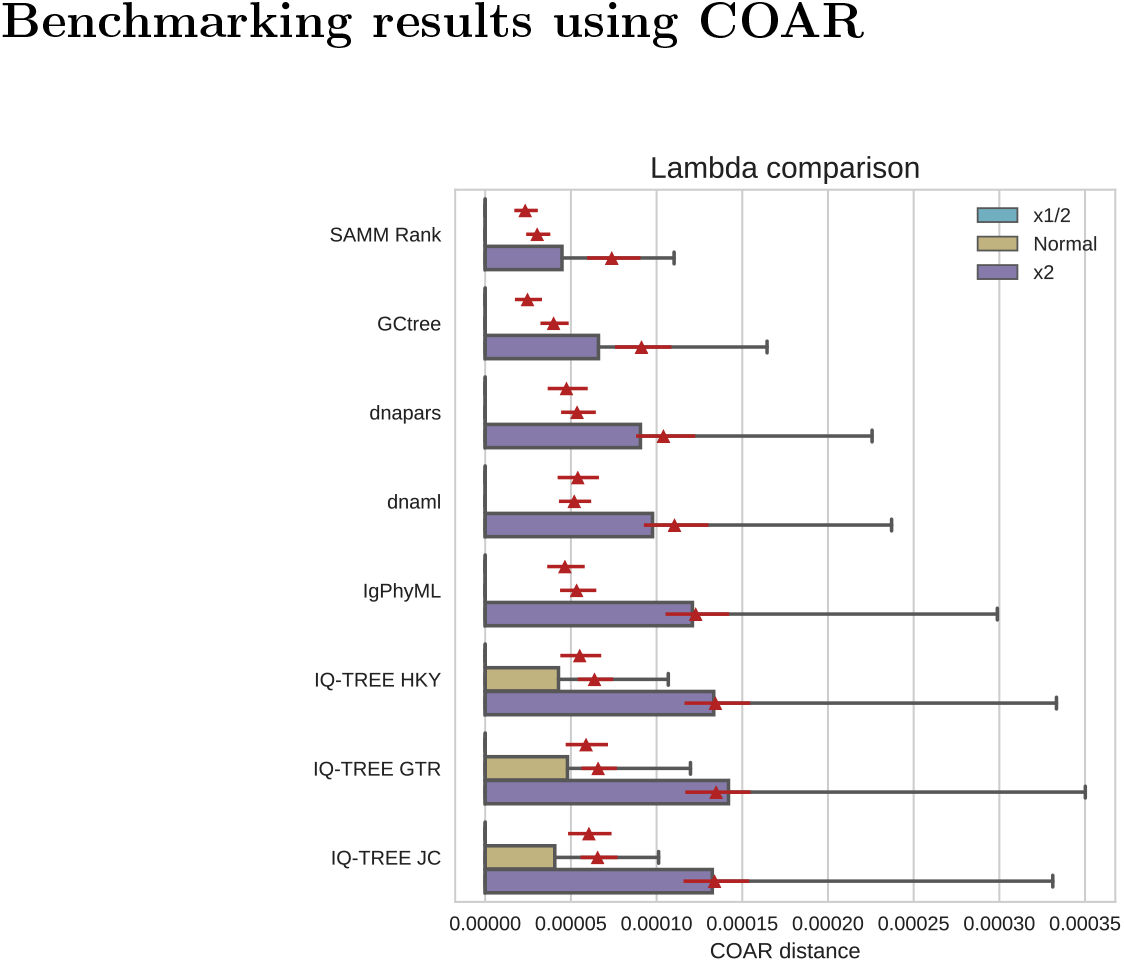
Neutral simulation showing COAR metric for mutation rates: “x1/2” = 0.1825, “Normal” = 0.365, and “x2” = 0.73.

**Figure S5:**
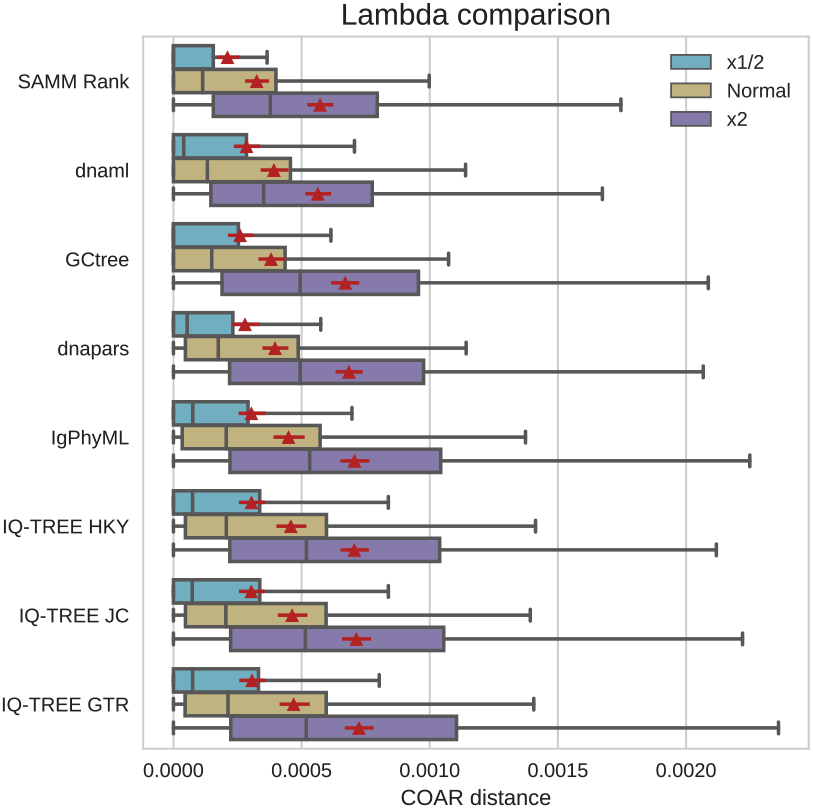
Affinity simulation showing COAR metric for mutation rates: “x1/2” = 0.1825, “Normal” = 0.365, and “x2” = 0.73.

**Figure S6:**
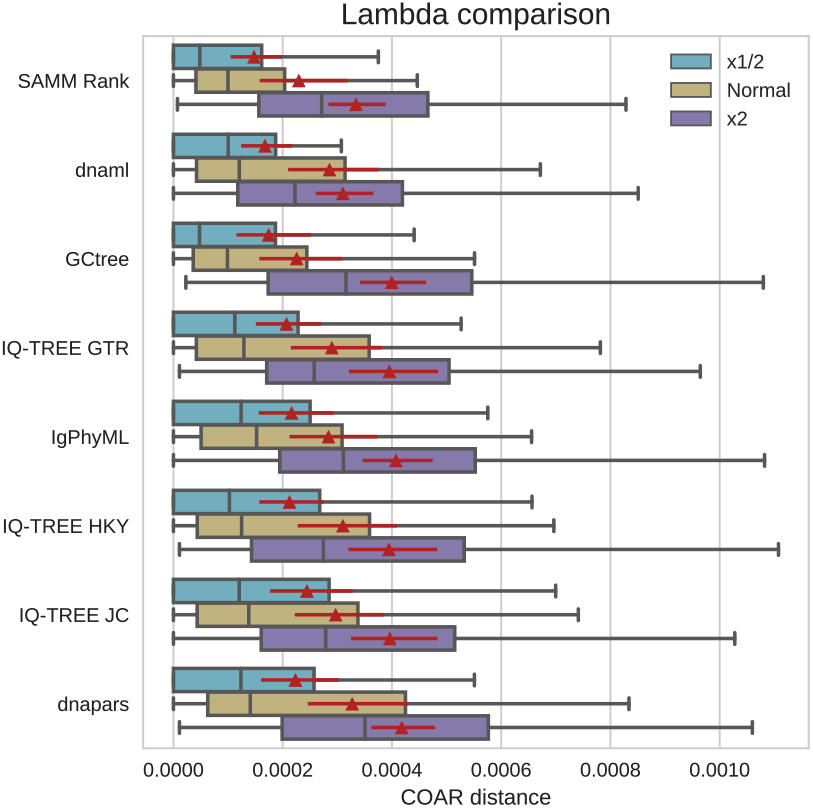
Affinity simulation with intermediate sampling (GC generation 15, 30 and 45) showing COAR metric for mutation rates: “x1/2” = 0.1825, “Normal” = 0.365, and “x2” = 0.73.

**Figure S7:**
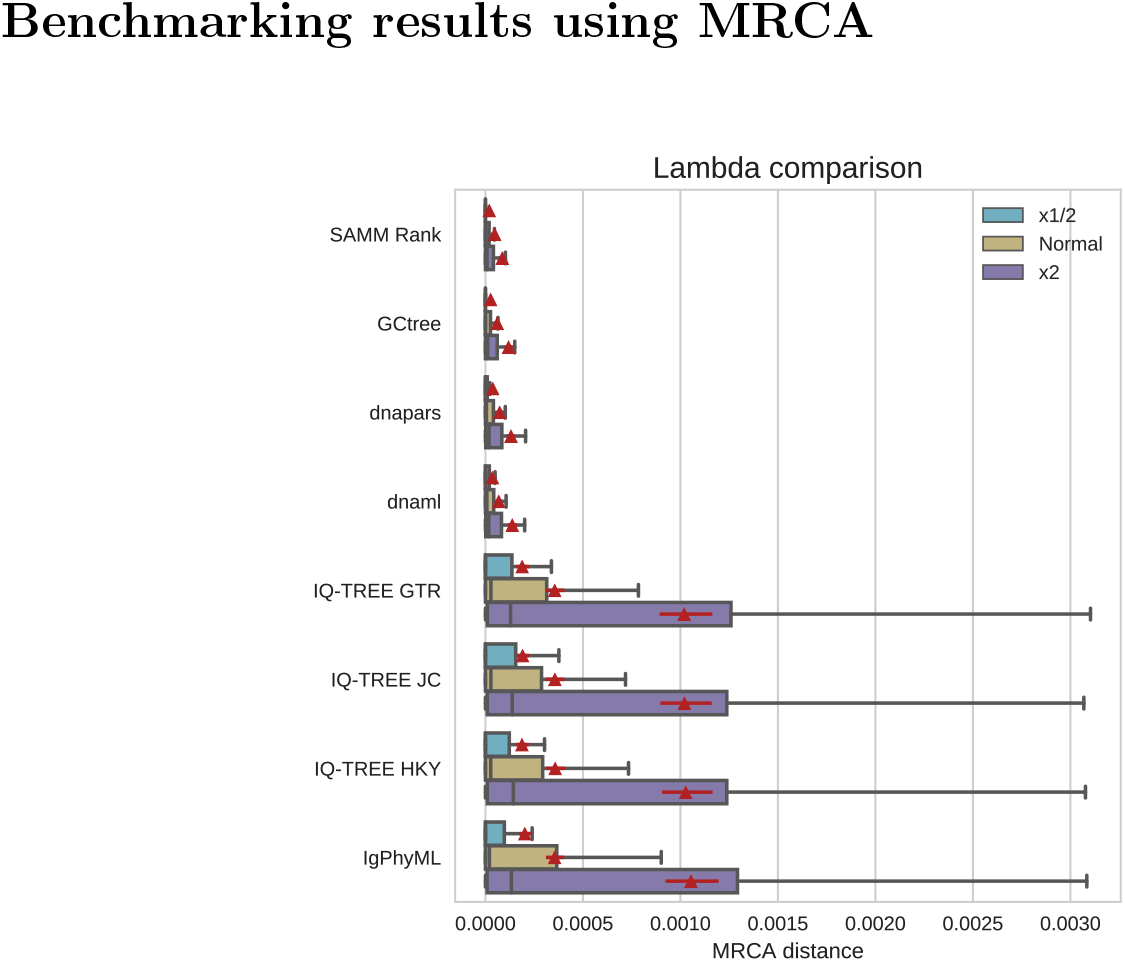
Neutral simulation showing MRCA metric for mutation rates: “x1/2” = 0.1825, “Normal” = 0.365, and “x2” = 0.73.

**Figure S8:**
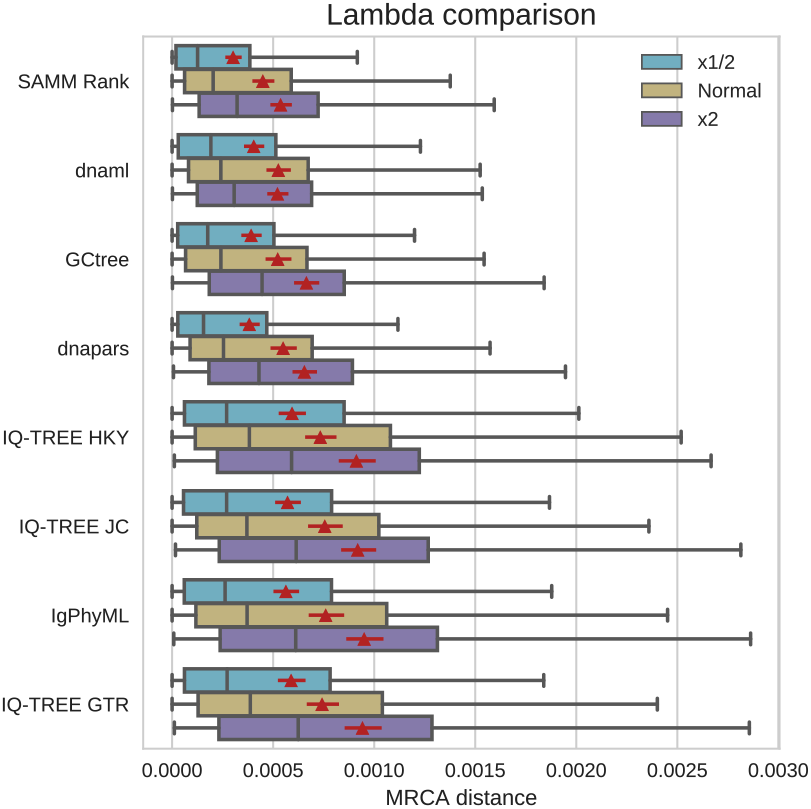
Affinity simulation showing MRCA metric for mutation rates: “x1/2” = 0.1825, “Normal” = 0.365, and “x2” = 0.73.

**Figure S9:**
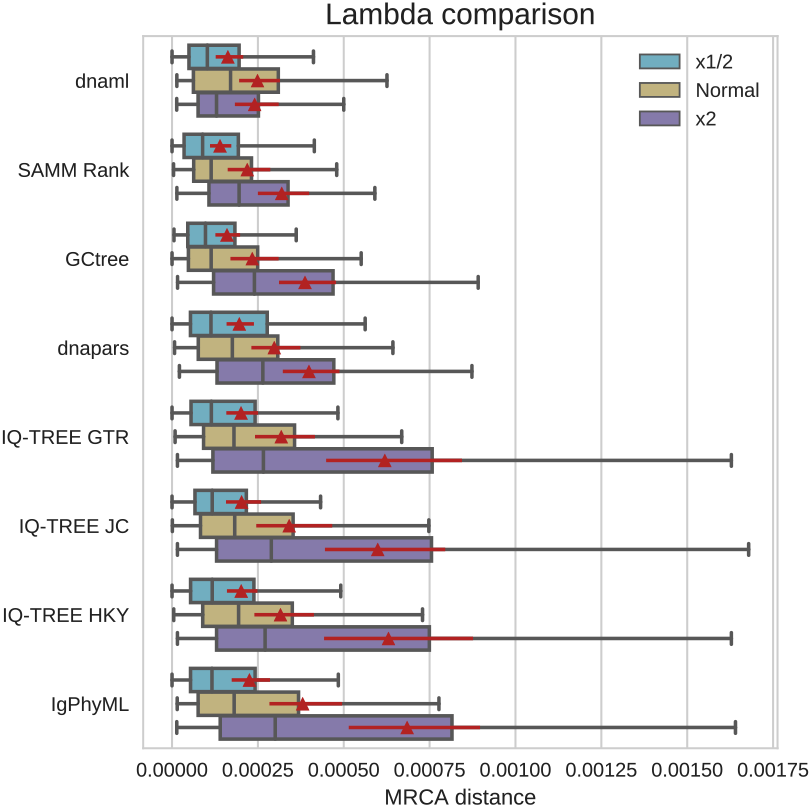
Affinity simulation with intermediate sampling (GC generation 15, 30 and 45) showing MRCA metric for mutation rates: “x1/2” = 0.1825, “Normal” = 0.365, and “x2” = 0.73.

**Figure S10:**
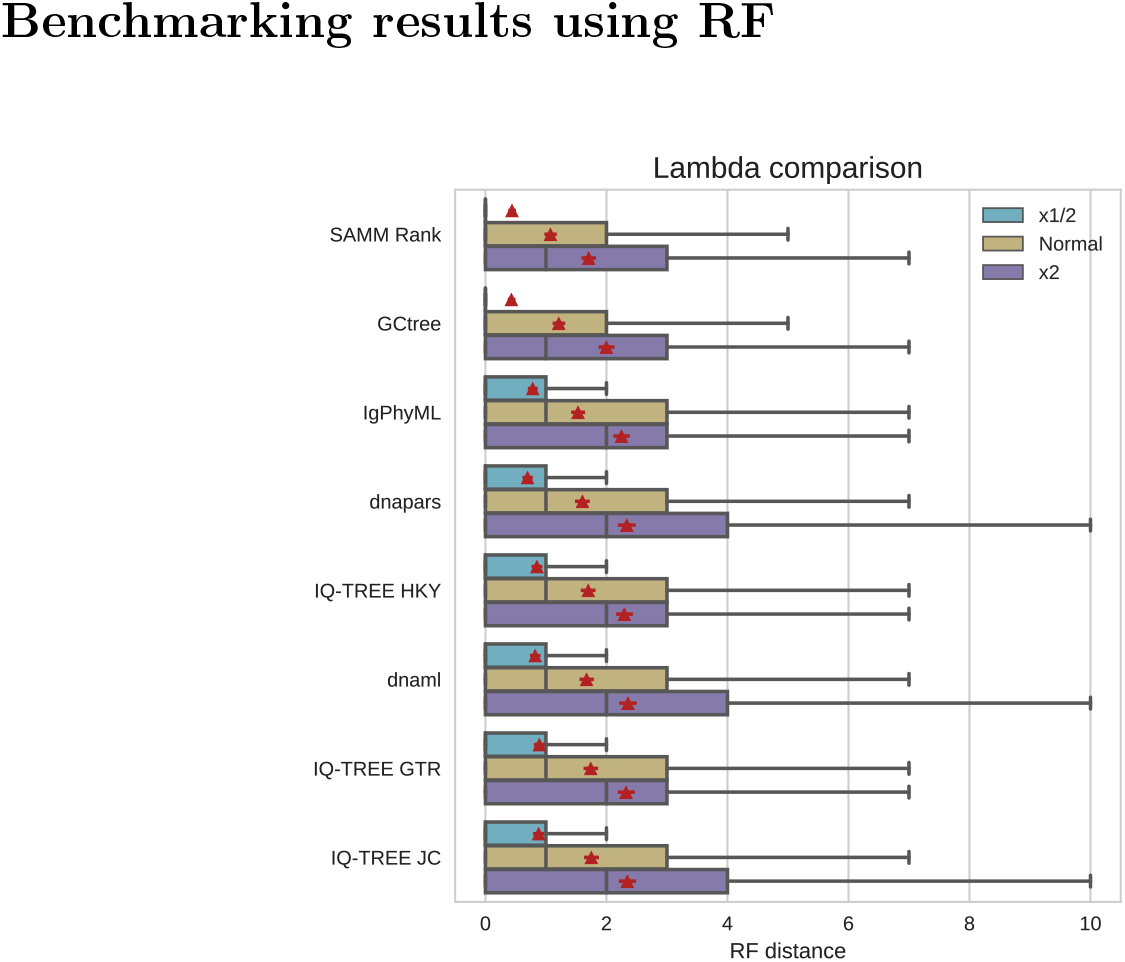
Neutral simulation showing RF metric for mutation rates: “x1/2” = 0.1825, “Normal” = 0.365, and “x2” = 0.73.

**Figure S11:**
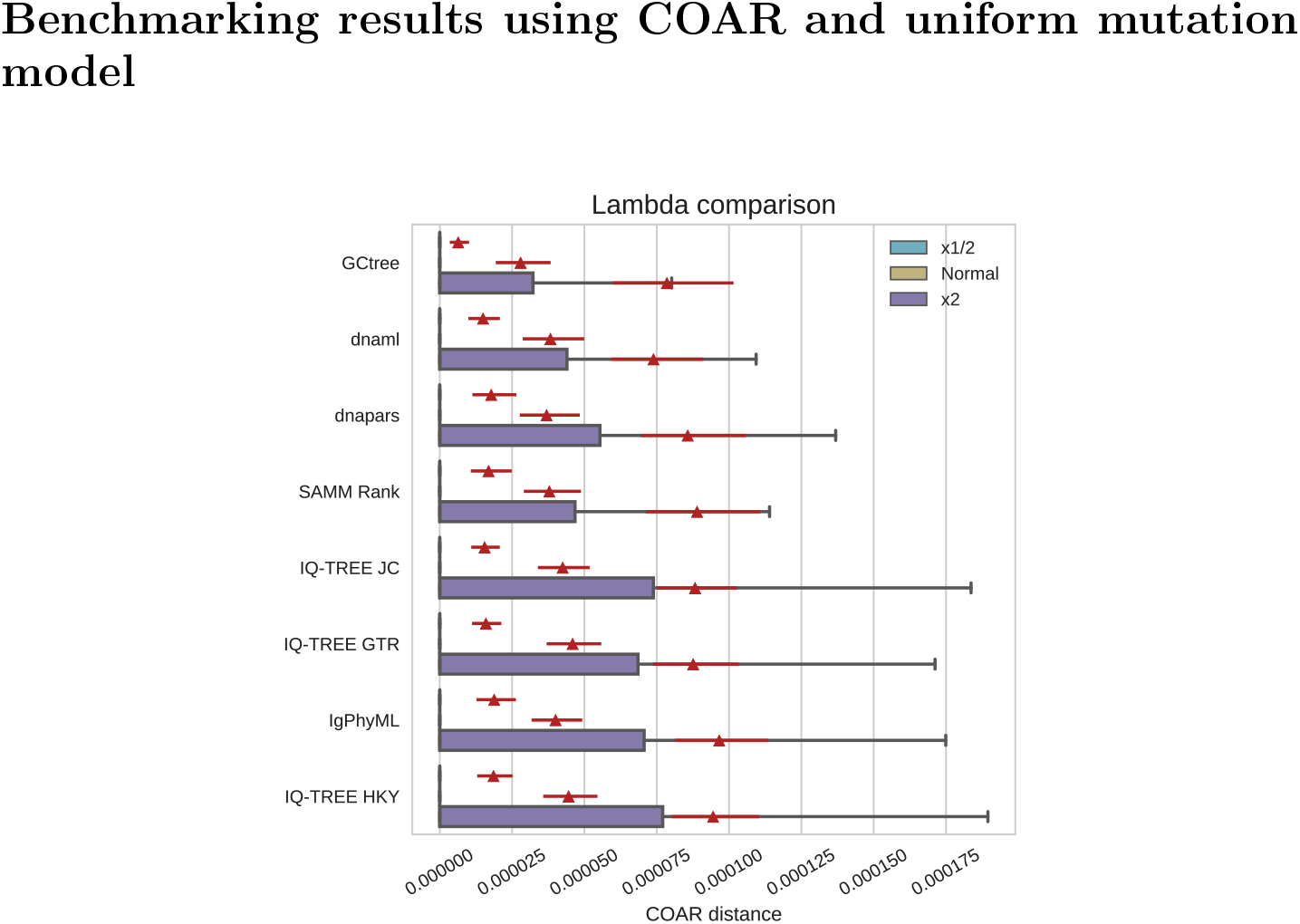
Neutral simulation showing COAR metric. Mutations were drawn from a uniform distribution over sites and substitutions using mutation rates: “x1/2” = 0.1825, “Normal” = 0.365, and “x2” = 0.73.

**Figure S12:**
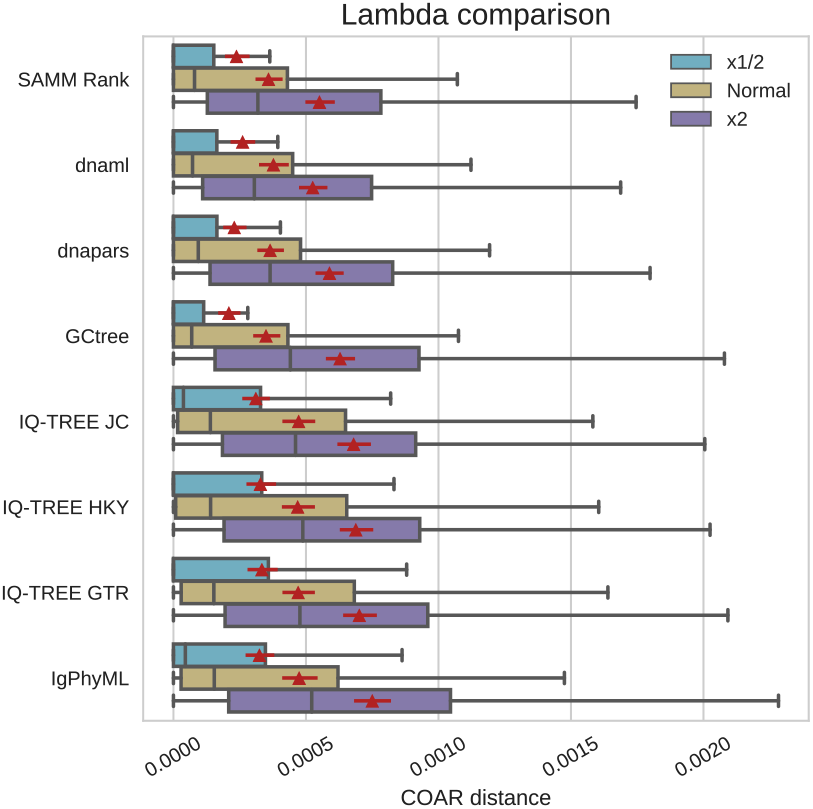
Affinity simulation showing COAR metric. Mutations were drawn from a uniform distribution over sites and substitutions using mutation rates: “x1/2” = 0.1825, “Normal” = 0.365, and “x2” = 0.73.

**Figure S13:**
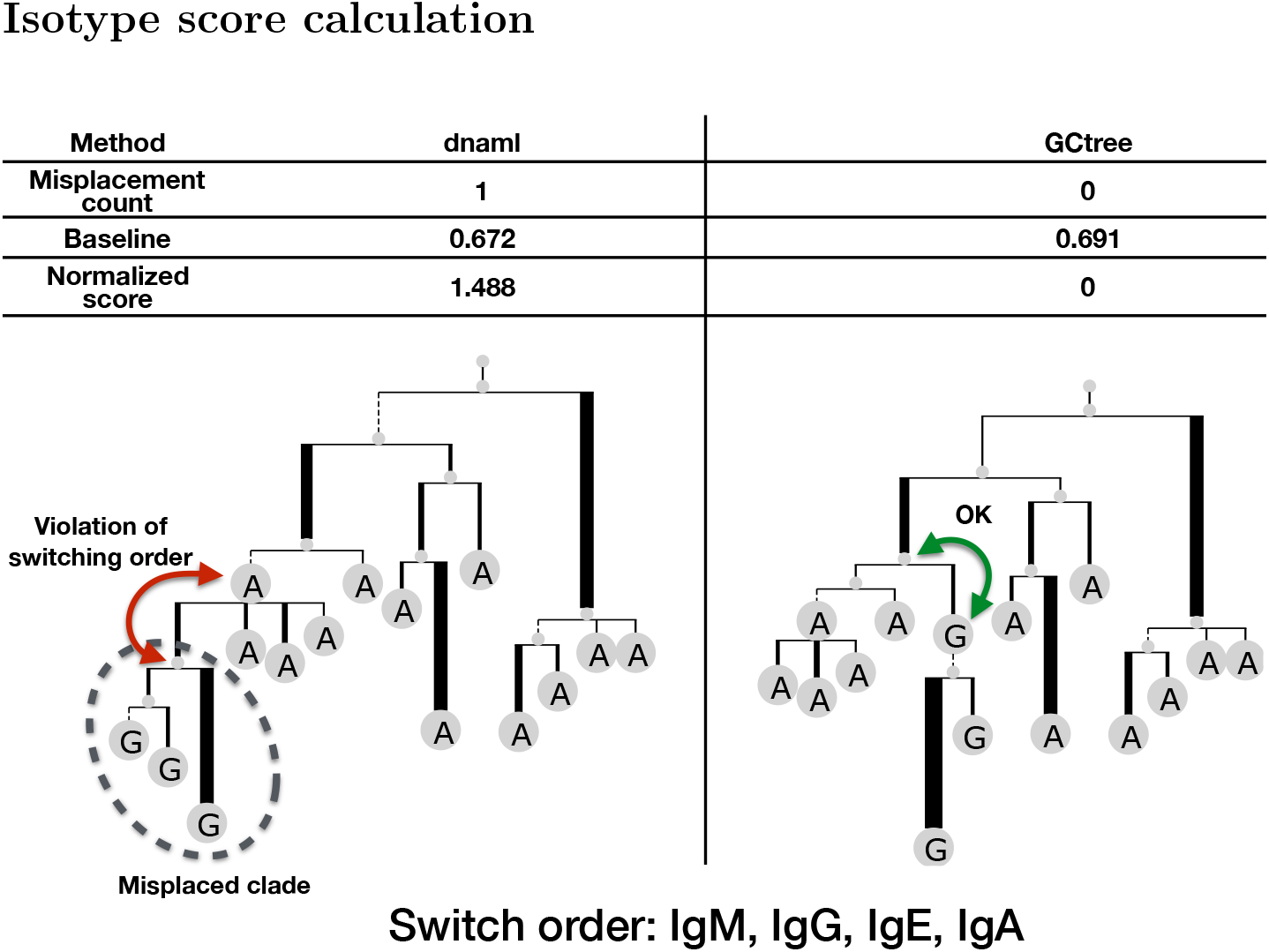
Example calculation of the isotype score. On the left: a tree inferred by dnaml where one clade has been misplaced resulting in a violation in the isotype switching order. One the right: a tree inferred by GCtree, on the same sequences, does not have any violations in the isotype switching order. The misplacement count is normalized by dividing it by a baseline score, found by taking the average misplacement score of 10,000 label shuffled trees of the same topology. The normalized score is also referred to as the “isotype score”.

### Isotype score comparison

The isotype score distribution was computed over 697 selected clonal families. The isotype score has a very high variance as can be observed in the 95% bootstrap confidence interval (10,000 replicates of sampling with replacement) of the mean. The comparison was run twice: once using the S5F motif model for SAMM ranking and another using SAMM’s own 5-mer motif model fitted on the mutations in the 697 selected clonal families. For all other tools these represent replicate runs. The replicated runs clearly exemplifies the uncertainty of the mean estimates e.g. IQ-TREE under the JC model was ranked 4th in the S5F replicate and 7th in the SAMM replicate. The only consistent feature is that IgPhyML ranks high (second best).

Since the data is paired a “significant” difference between two compared tools should be calculated using a paired statistical test. One way of showing the comparison between paired data is to subtract it pairwise and compute confidence intervals on the differences. We do this by treating the best tool as a reference point and finding the distribution of differences to other tools. From the 95% confidence intervals of these differences one can accept/reject the null hypothesis of no difference in the means. This hypothesis test shows that both the SAMM and GCtree parsimony ranking strategies are significantly better than dnapars.

**Figure S14:**
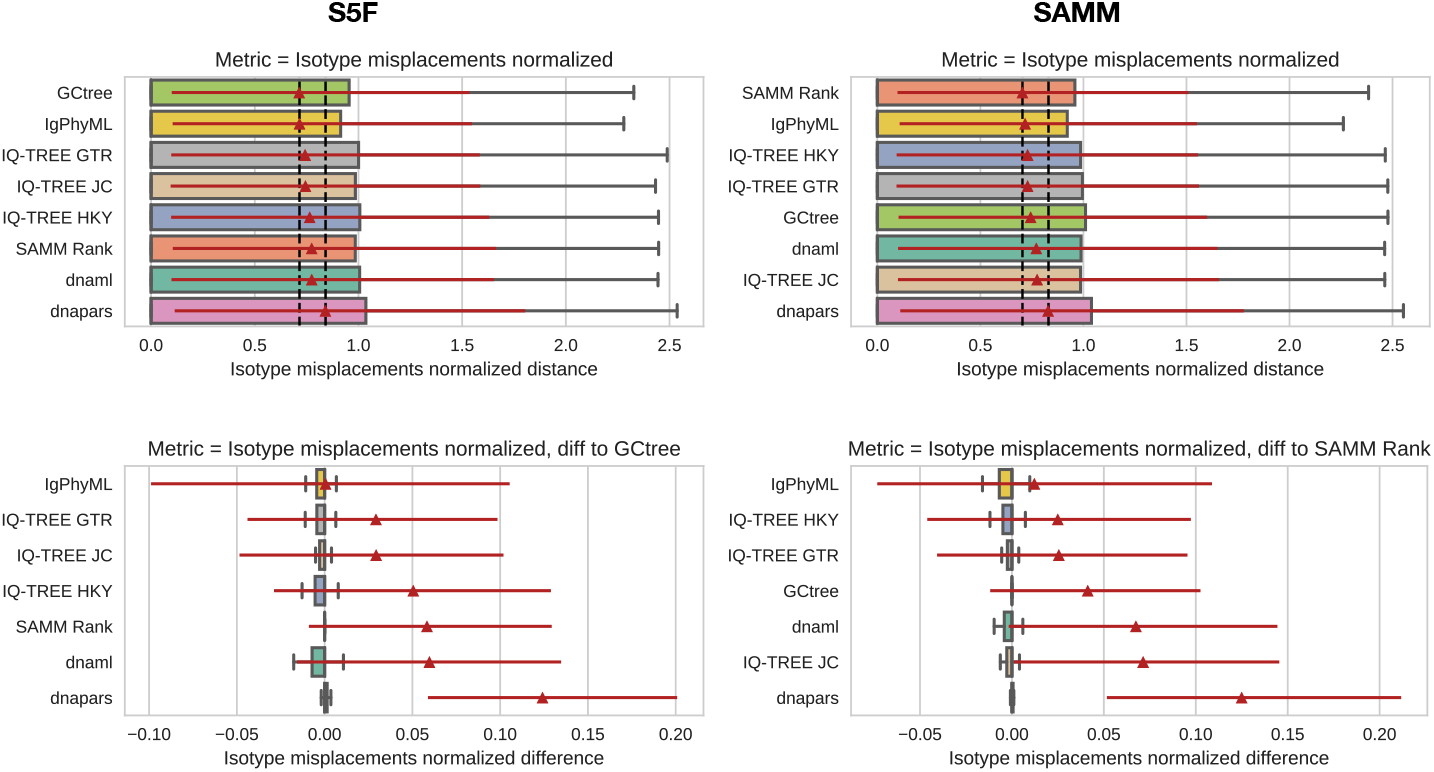
Isotype score distribution on 697 selected clonal families. In the left column: SAMM ranking uses the S5F model. In the right column: SAMM ranking uses its own motif model fitted on the mutations in the input data. Upper row shows the isotype score distribution, lower row shows the isotype score distributions after subtracting the isotype scores of the best ranked tool.

### Simulating affinity maturation

In this section we will describe our BCR sequence simulation framework in depth, first by introducing the neutral process which is the foundation of all our simulations, and then going on to motivate and derive a model that incorporates BCR affinity and antigen competition to define sequence fitness.

#### Neutral model

The neutral process can be viewed as a model of cell divisions, where at each cycle through the GC a cell can either die or produce a number of offspring, and each offspring has some probability of carrying mutations. Offspring numbers larger than two are used to approximate multiple cell divisions in a single GC cycle. The root sequence (naive BCR) is given at the simulation initialization as a starting point from where the tree is evolved until the simulation is stopped. Cell division is controlled by a Pois(λ) progeny distribution, and at each GC cycle all progeny cells will undergo a mutation process. The number of nucleotides to mutate is drawn from another Poisson distribution (Pois(λ_mut_)) and introduced sequentially into the sequence using a substitution model. Sequential introduction of mutations allows the possibility of back mutations. We use the S5F mutation model (17) to introduce mutations, which describes mutability and substitution preferences of the middle base of all 5-mer DNA motifs. However, a 5-mer mutability cannot be used directly on sites at the start or end of a sequence because of missing context, therefore we fill in missing context with the unknown base, N, and average over all possible motifs fitting into this ambiguous context.

Termination of the neutral branching process is achieved in either of three ways: 1) by simulating under a subcritical process (λ < 1) (39) and following it until extinction, 2) by using a stopping time *T*, or 3) by stopping after a population of *N* cells has been reached. Sequences are then sampled from the tree leaves. In addition we introduced a parameter for down-sampling the cell population to *n* cells. Model parameters are tabulated in Table S1.

**Table S1:** Parameters used in the neutral simulation.

| Parameter | Description |
| --- | --- |
| $\lambda$ | $\text{Pois}(\lambda)$ progeny distribution |
| $\lambda_{\text{mut}}$ | $\text{Pois}(\lambda_{\text{mut}})$ mutation distribution |
| $T$ | Stopping time |
| $N$ | Stopping number of sequences |
| $n$ | Down-sampled number of sequences |

#### Simulations with affinity selection

To model the affinity maturation process with selection we will use the exact same framework as described for the neutral process, but now the Pois(λ) progeny distribution is no longer constant. We consider the magnitude of λ as the fitness of a cell. In the neutral model λ is a fixed constant resulting in a completely flat fitness landscape, as opposed to a system with selection where the fitness landscape is a more complex and rugged surface. The following subsections will describe a model with the simple purpose of defining a function to calculate the fitness of any BCR sequence. The fitness is measured in terms of a single λ^(i)^ associated to each cell and defining a cell specific progeny distribution. Thus, selection is condensed into a dynamic λ, and this is the only difference to the neutral process.

#### Model concept and biological assumptions

Let us make some basic assumptions to keep later definitions simpler. First, the system we intend to model is the affinity maturation process happening in the GC, assumed to be driven by the BCR’s affinity towards a single target antigen. A real GC reaction is seeded by 50-200 naive B cells, however, due to the extensive competition they often completely “resolve” in later stages of affinity maturation, resulting in cells with only a single common naive B cell ancestor i.e. monoclonallity (20). We do not attempt to model this inter-clonal competition so we make the simplifying assumption that the simulated GC is seeded by a single naive B cell. In our model it is the BCR amino acid sequence that is under selection, thus we ignore the possible fitness effects of synonymous mutations.

The GC is modeled with constant volume and constant total concentration of antigen. B cells compete for this limited antigen. B cells with high affinity BCRs will bind more antigen and are more likely to undergo cell division and vice versa for low affinity BCRs. Binding equilibrium is assumed to be instantaneous and the progeny distribution for a B cell is evaluated as a function of the BCR occupancy at this equilibrium. Affinity is a function of the BCR sequence and its amino acid sequence distance from the best BCR (here called the mature sequence). Once a new cell has been created this changes the binding equilibrium which then needs to be updated. A GC cycle in the simulation is defined by one iteration through all the cells to evaluates their progeny distributions. Cartoon overview in Figure S15.

#### Kinetic model of BCRs binding antigen

In the following we derive the fraction of a B cell’s BCRs bound to the antigen in a GC (BCR occupancy). This is then extended to a situation with multiple B cells with different BCR affinities.

First, consider the BCRs of a single B cell as free molecules with a total concentration of [*B*_total_], then the BCR occupancy at equilibrium is:

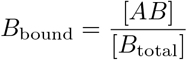

Where [*AB*] is the concentration of BCRs bound to antigen. We need to derive a solution to calculate *B*_bound_.

The binding equilibrium between free antigen ([*A*]), free BCRs ([*B*]) and BCR bound antigen ([*AB*]) is:

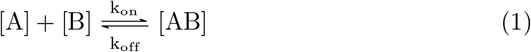

**Figure S15:**
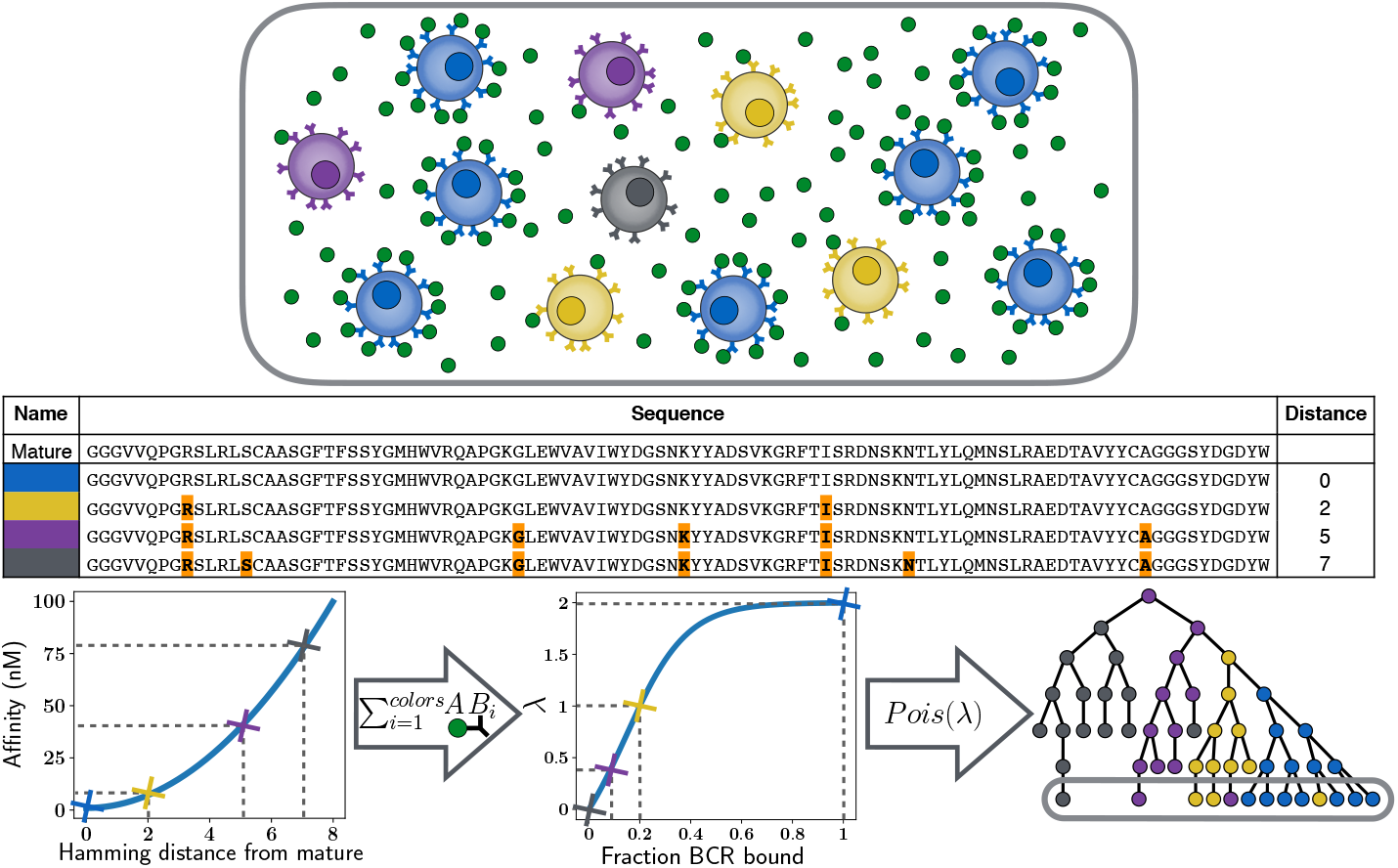
Simulation overview. The system is considered as a closed environment with free floating antigen and a number of B cells presenting BCRs on their surface, (top panel). Different colors correspond to different affinity BCR sequences. In the middle panel a sequence alignment shows the distance between BCR sequences and the mature BCR. Bottom panel shows first how distance from the mature BCR is converted to affinity, then how the fraction of bound BCRs is transformed to a λ defining the progeny distribution. Rightmost of the bottom panel shows the lineage tree with an ellipse marking the B cells of the current generation also displayed in the top panel.

The on- and off-rate of binding is expressed as constants *k*_on_ and *k*_off_. Affinity can then be expressed as:

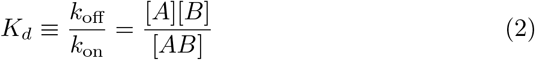

Isolating [*AB*]:

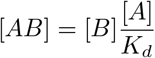

Substituting [*B*] for its expression from mass conservation, [*B*_total_] = [*B*] + [*AB*]:

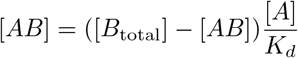

Which rearranges to the result:

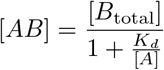

Then extending the model for binding equilibrium of a single BCR sequence to one with multiple BCR sequences just requires indexing:

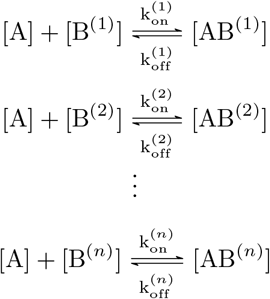

The same solution applied and because all B cells compete for the same antigen, each [*AB^(i)^*] is dependent through the concentration of unbound antigen:

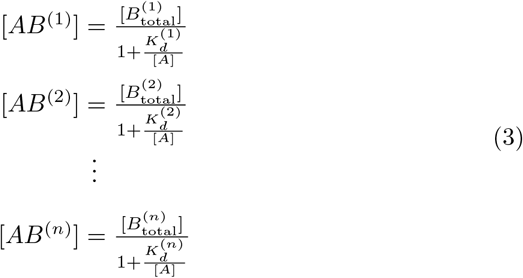

Now introducing mass conservation for the antigen *A*:

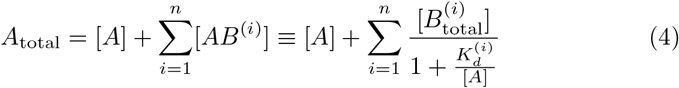

By rearranging to a polynomial form the system can be solved by root finding to calculate [*A*] which is then used to find all the [*AB^(i)^*]’s and transformed them to 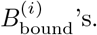

This is a solution to a model of BCR competition in the GC but to make this work we also need a definition of BCR affinity as well as a way of transforming BCR occupancy to fitness in the sequence simulation.

#### Defining affinity for a sequence

Here we describe how to define the affinity 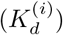 of each BCR. A numerical affinity value can be generated by transforming a BCR sequence (*S^(i)^*) into a number that represents affinity. Formally, this would be a function: 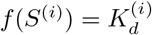. Consider that the BCRs in a GC are evolving towards a specific mature sequence, denoted *S^M^*. A mature sequence is the sequence with the highest affinity and fitness. We will define a fitness landscape around this mature sequence using Hamming distance between amino acid sequences: *d_H_*(·, ·).

Let us define the affinity of the naive input sequence as 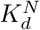 and correspondingly the affinity for the mature sequence as 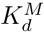. Now, we can define an arbitrary function with reference points in 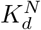 and 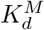, that transforms a distance between *S*^(i)^ and *S^M^* to an affinity:

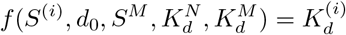

Where *d*_0_ = *d_H_* (*S^N^,S^M^*) is the distance between the naive and mature sequences. There are two constraints we want to impose. If the BCR sequence is: 1) equal to the naive sequence (*S^N^*) it takes the affinity of the naive BCR 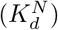, and 2) equal to the mature sequence (*S^M^*) it takes the affinity of the mature BCR 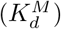:

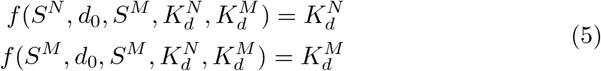

A flexible function for transforming distance to affinity is the family of power transformations which we define with the two conditions satisfied as:

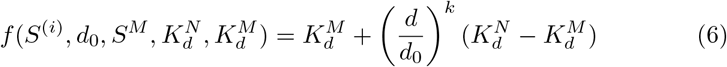

Where *d* = *d_H_*(*S^(i)^,S^M^*) is the distance between the input and mature sequences. The exponent, *k*, can be chosen to adjust the mapping between distance and affinity, with the restriction that 0 < *k*because the function < ∞ (Figure S16).

**Figure S16:**
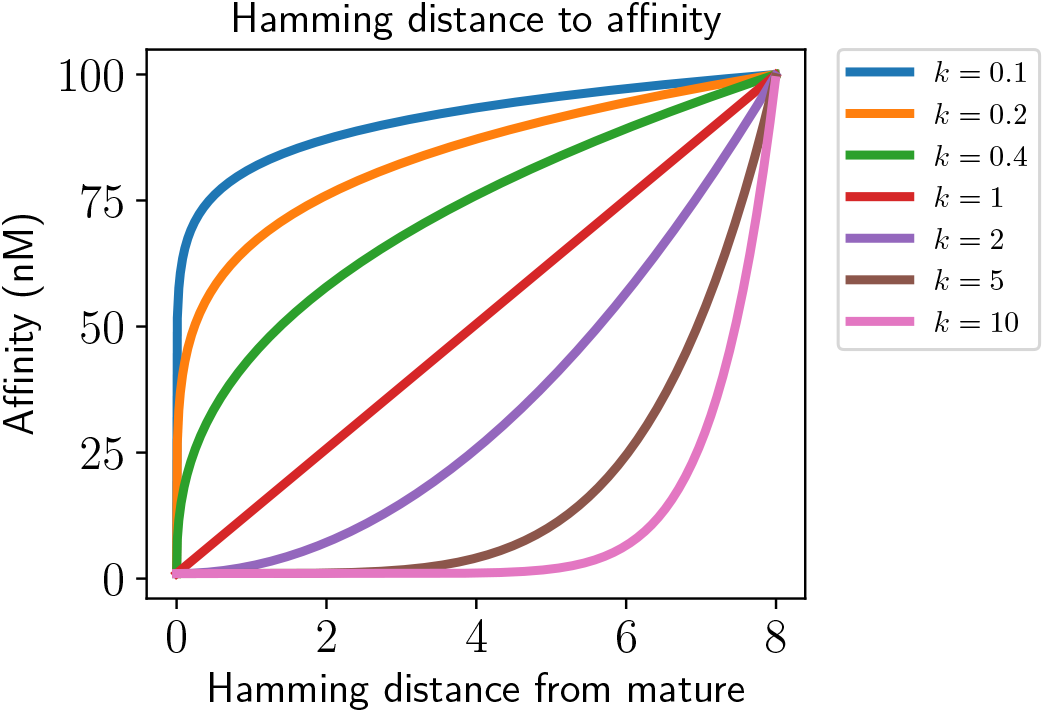
Varying the exponent *k* in (6) to achieve different mappings between distance and affinity. Naive and mature affinity is held constant, 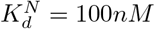 and 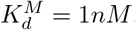.

In a real affinity maturation process there may be many different BCR sequences that are practically equally fit e.g. this will happen when multiple amino acids are equally fit on a given position, and it will also happen if there are multiple distinct maturation paths that end up with equally fit BCRs. Our model deals with this by allowing multiple mature sequences and then determining the affinity based on the shortest distance to any of these:

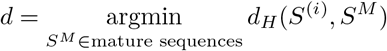

#### Transforming BCR occupancy to fitness

Equipped with a sequence to affinity mapping and a method to solve the binding equilibrium in a population of BCRs the last element necessary is to couple BCR occupancy to fitness. This is achieved through the progeny distribution; if 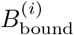 is small the progeny distribution should favor terminating the B cell and opposite, if 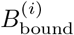 is large the progeny distribution should favor cell division. The Poisson distribution will reflect this behavior by setting λ^(*i*)^ small when 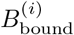 is small and λ^(*i*)^ large when 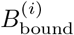 is large. However, it is unrealistic that there should be a one-to-one mapping between 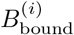 and λ^(*i*)^ and therefore we need a function for transformation: 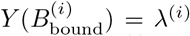. The function should allow specification of lower and upper bounds on λ^(*i*)^, a threshold (*f_full_*) on 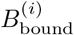 when more bound antigen does not have any fitness effects (Figure S17) and another threshold 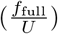 defining 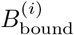 when the progeny distribution transitions between a subcritical and a supercritical process (λ^(*i*)^ = 1) (39) (Figure S18). These requirements can be accommodated by the generalized logistic function:

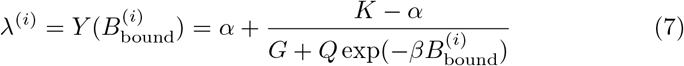

*G* is chosen to be the typical logistic function value of 1. *K* is the upper bound on λ^(*i*)^ and is set to 2 (slightly larger than the λ = 1.5 fitted for the neutral branching process). *α*, *β* and *Q* are found using three conditions:

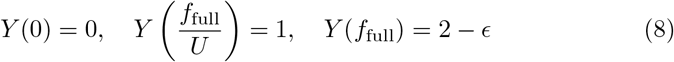

The solution is undefined in *Y*(*f*_full_) = 2 because the function is asymptotically growing towards 2, therefore e can be regarded as a small value (e.g. 10^-3^) so that *Y*(*f*_full_) ≈ 2. The constant *U* in condition 2 can be adjusted to set the value of 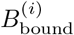 resulting in λ^(*i*)^ = 1. Using these conditions *α*, *β* and *Q* can be found and the logistic function is fully defined. *α* can be interpreted as the lower asymptote of the function. *β* is the steepness of the function and it is coupled to the *Q* parameter and follows it according to the three conditions in (8).

#### Parameter choices

We define the maximum fitness to be attained at 100% BCR binding, hence we fix *f*_full_ = 1. The infliction point parameter *U* is chosen to reflect our expectation that initially, when only a few BCRs are bound and stimulation is low, there will be a linear increase of the stimulus when antigen binding increase, and at some point close to *f*_full_ the increase in stimulus levels out. This expected shape is recapitulated by a choosing *U* = 5 (Figure S18).

**Figure S17:**
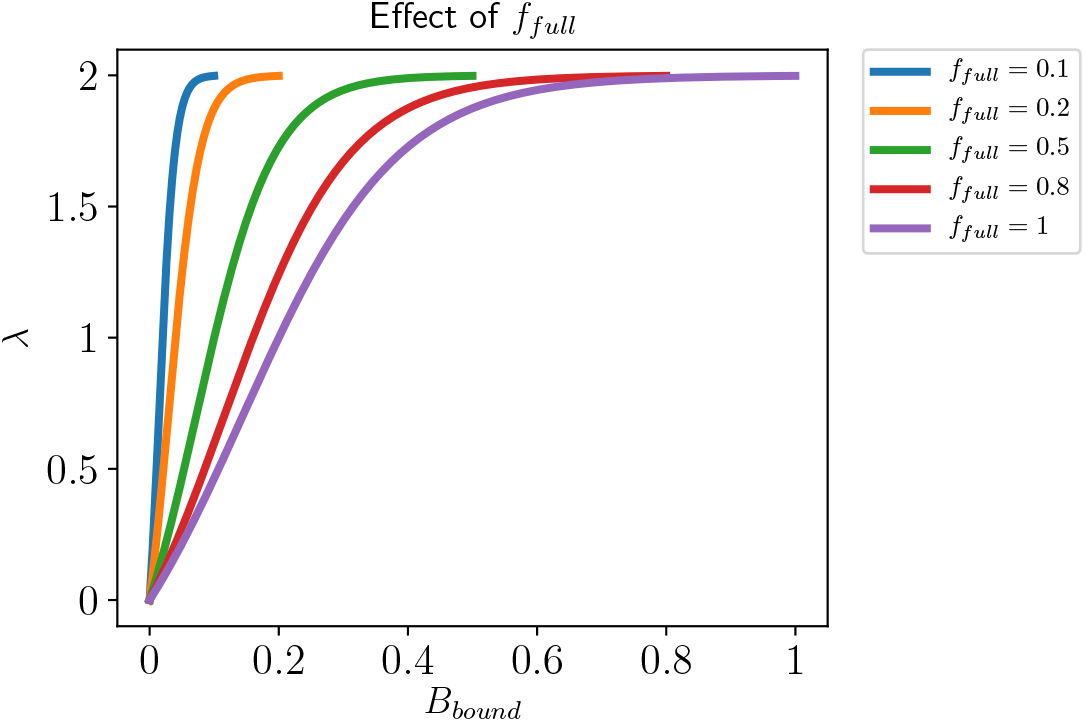
Using a constant U = 5, changing the *f*_full_ parameter in the conditions in (8) to change the point where *B_bound_* reaches the λ plateau.

**Figure S18:**
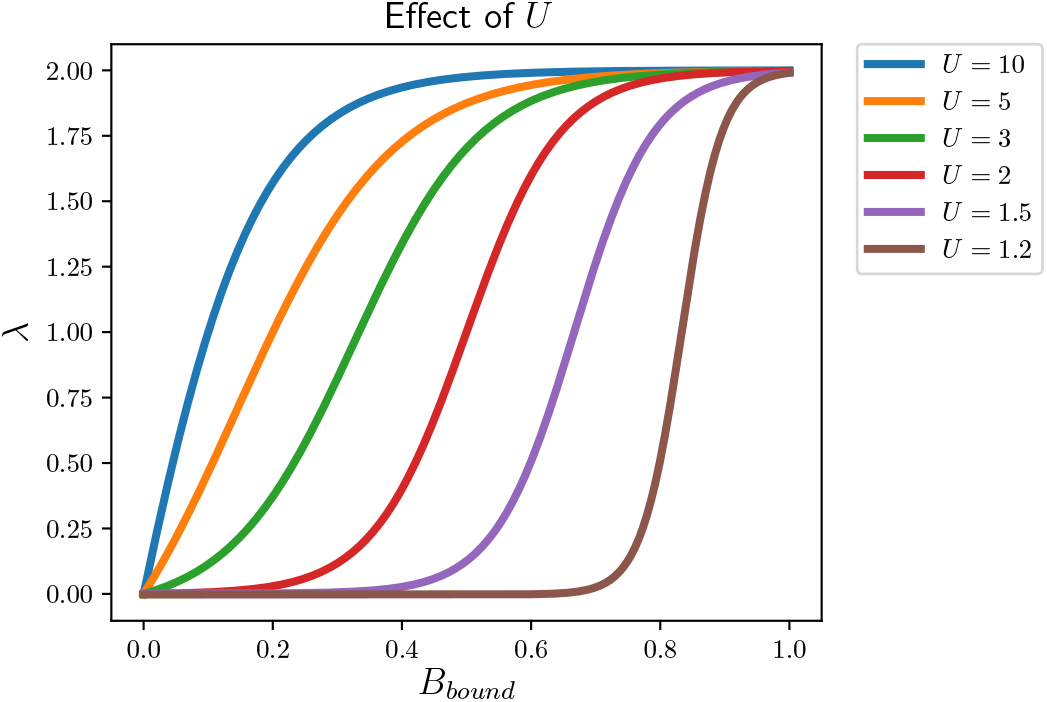
Using a constant *f*_full_ = 1, changing the *U* parameter in the conditions in (8) to achieve a shift of the inflection point at λ = 1 on the *B_bound_* axis.

The total concentration of antigen (*A*_total_) needs to be defined to solve the binding equilibrium. To do this we need to introduce the concept of a carrying capacity of the simulated GC, which is defined as the number cells a GC is able to support in its micro environment. The carrying capacity is determined mainly by the total concentration of antigen since binding to antigen controls the progeny distribution. BCR affinity is also influencing antigen binding and therefore must influence the carrying capacity, but at high affinity nearly all antigens are bound and hence the total antigen concentration is the most influential determinant of GC carrying capacity. At Pois(1) the progeny distribution is only just sustaining the population size of the GC, and given condition 2 in (8) this happens at 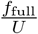. Then, under the assumption that the population of B cells all have identical BCR sequences, the maximum carrying capacity is:

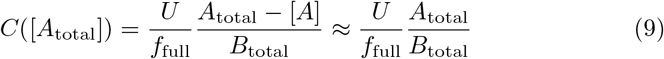

Using a carrying capacity of 1000 (41, 74) we can calculate *A*_total_. We note that simulations are generally robust to different parameter choices (Figure S19).

**Figure S19:**
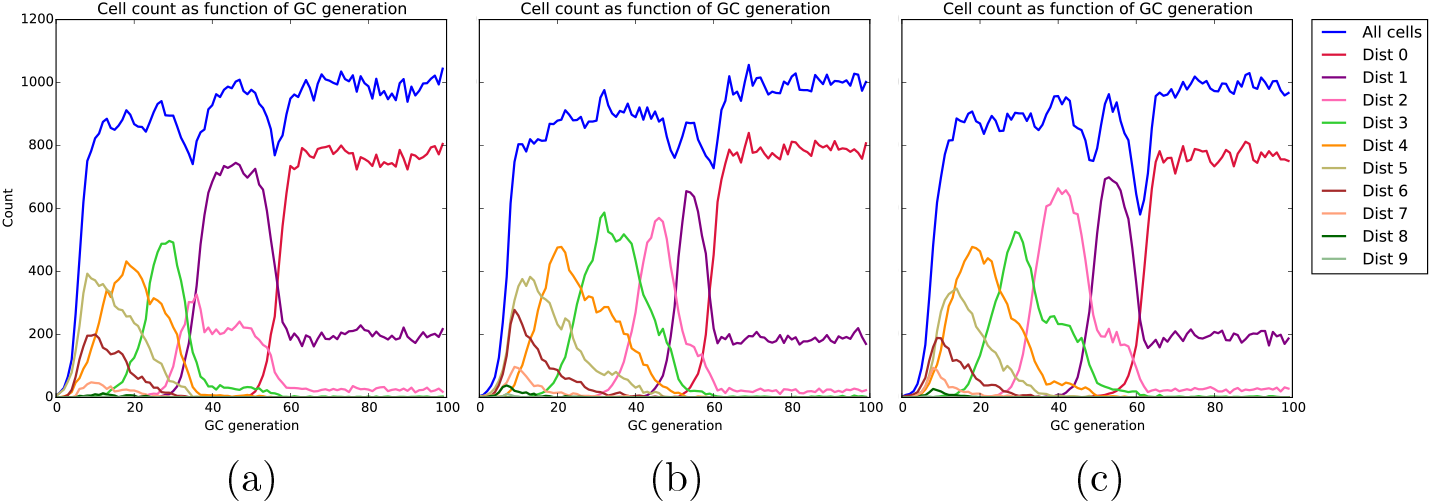
Simulation with affinity selection for varying magnitudes of *f*_full_. (a) *f*_full_ = 1, (b) *f*_full_ = 0.5 and (c) *f*_full_ = 0.05. Simulations with *d*_0_ = 10, *U* = 5 and [*A*_total_] adjusted to obtain a carrying capacity of 1000 cells. Each simulation was run for 100 generations and the composition of sequence distances to their closest mature sequence are plotted for each generation.

In the transformation from distance to affinity in (6), we have to make a choice about which exponent to use. We would like to disallow sequences drifting far away from the mature sequence by enforcing a positive exponent. Furthermore, we require that each Hamming distance step between the naive and mature sequences has a substantial affinity effect, and therefore *k* = 2 is used.

The amino acid sequence distance between the naive and mature sequences, do, is set to 5. The *K_d_* for a naive sequence is likely in the low micro molar range range of 10^-6^ − 10^-7^*M*, while the mature affinity is in the nano or subnano molar range of 10^-8^ − 10^-10^*M* (75–78) (*M* is used to denote molar concentration). We choose the naive sequence to be 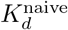 = 10^-7^*M* (100*nM*) and the mature to be 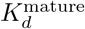 = 10^-9^*M* (1*nM*), giving a large span in affinity to select on. Based on approximating the GC as spheric, and using the experimental data for average GC diameter and BCRs per B cell, the model is fully defined in nanomolar concentrations. All necessary constants are tabulated in Table S2.

**Table S2:**
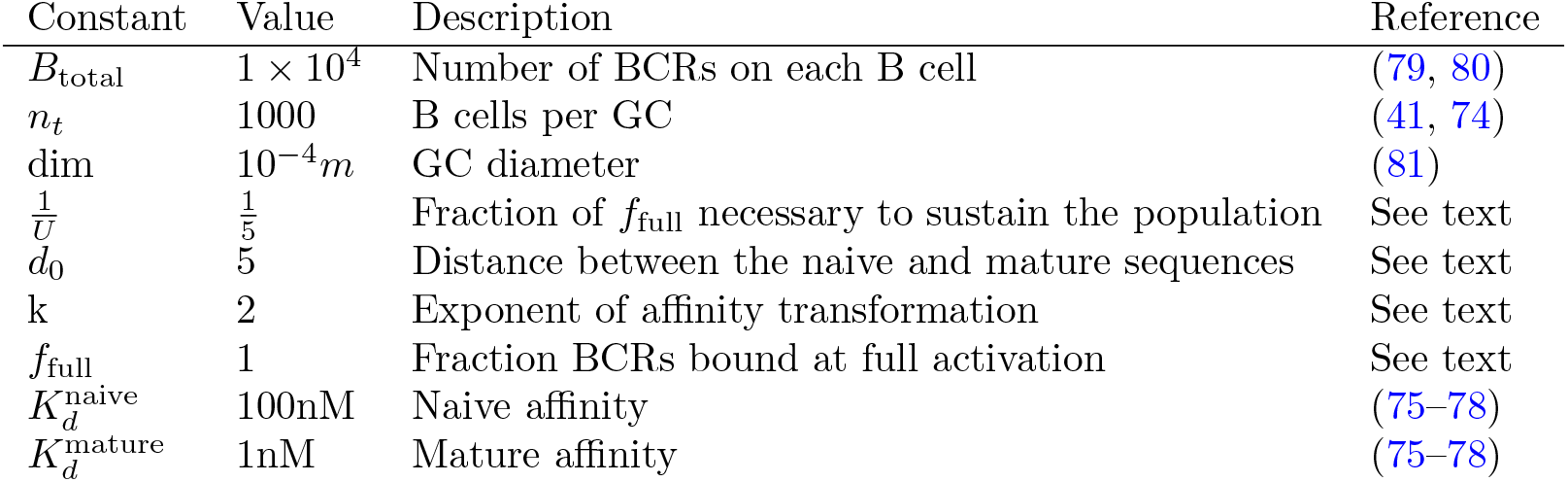
Constants used in the model of affinity simulation.

#### Correctness of ancestral reconstruction

In the following section we will introduce a benchmark metric for ancestral sequence reconstruction, which we call “correctness of ancestral reconstruction” (COAR). The correctness of a reconstruction compared to the true evolutionary history can be measured by multiple similarity measures e.g. topological similarity, branch length similarity and sequence similarity between inferred and real ancestors. All these measures are inter-dependent e.g. the inferred sequences are affected by the branch lengths and the topology and the branch lengths are conditioned on a topology etc. And while inferring correct tree topology is important in its own right, the correctness of the inferred ancestral sequences are the foremost important objective of most BCR phylogenies when these sequences are used for applications involving DNA synthesis, protein expression and functional testing. For this reason, the sole purpose of the COAR metric is to capture the correctness of the inferred ancestral sequences. In particular, we would like to propose a loss function that does not penalize a phylogeny when minor parts of the tree topology is incorrect while ancestral sequence reconstruction is perfect.

The purpose of COAR is to compare two trees built with the same leaves; let us call these the true and inferred tree. When performing ancestral sequence reconstruction the desired result is often to reconstruct the internal nodes in the direct path going from a leaf to the root, as illustrated in Figure S20. This path is extracted by starting at a leaf node and traversing upwards, parent by parent, until the root is reached. In the following, this list of sequences will be referred to as the ancestral lineage. The correct ancestral lineage is the objective of COAR, and we construct the COAR value so it represents the expected per-site error in such a reconstruction. Following the example in Figure S20, often there will be small differences in tree topology between the true and inferred trees, and these will likely make the number of internal states in the ancestral lineages differ. This makes comparison difficult because two lists of different length cannot be element-wise compared. The lists could be made equal length by adding gaps, but then a systematic way of adding these would be necessary.

The basis of COAR is a list comparison progressing element-wise through the list i.e. element 1 in list 1 compared to element 1 in list 2, next, element 2 in list 1 compared to element 2 in list 2 etc. For lists of similar length the list comparison is easy, it will simply be the cumulated distance from list element comparisons, corresponding to the sum of Hamming distances between inferred and true ancestors in the lists. When lists are not equally long, one or more gaps must be introduced into one of the lists; we choose to do so in such a way that the list similarity is maximized. This is an alignment problem with matches/mismatches/gaps and it can be efficiently solved using the Needleman-Wunsch algorithm (82). We define it as a global alignment so that it has to start at the root and end at the leaf because both states are known for the true and inferred phylogenies. We further restrict the Needleman-Wunsch algorithm so that gaps are only allowed to be introduced into the shortest of the two lists being aligned, this forces the maximum number of node comparisons.

One interpretation of the COAR value is that it is the distance between the true and inferred mutation histories, as illustrated in in Figure S21. In this representation of an ancestral lineage the root and the leaf are two fixed states with a continuous mutation process running between them. The internal nodes

**Figure S20:**
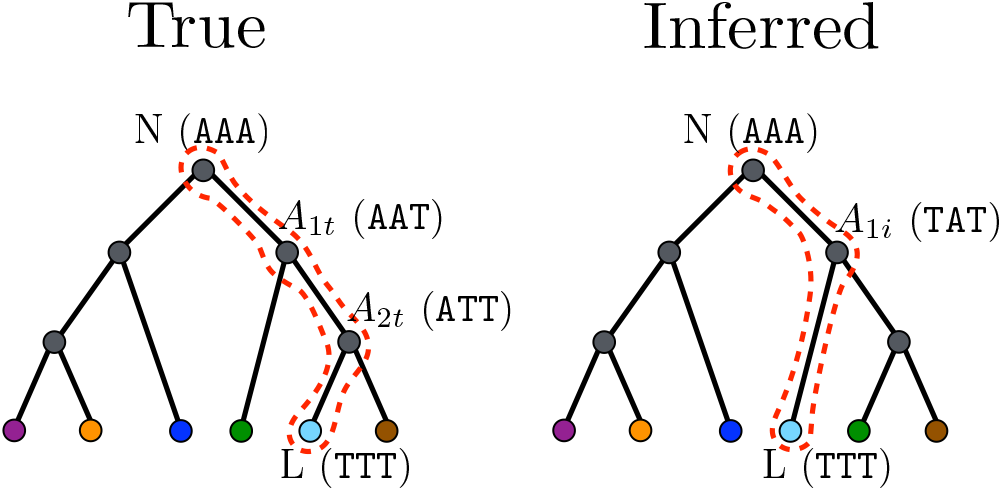
True vs. inferred tree with colored leaves and grey ancestral states. Reconstruction from the light blue leaf is marked by a dashed red line and annotated with genotypes in parenthesis. N is the naive sequence, L is the leaf sequence and the As are ancestors 1, 2, …, *n* with either true or inferred marked by *t* or *i*, respectively, appended to the subscript. The inferred tree has misplaced the branch leading to the light blue node, resulting in a missing ancestral sequence.

in the ancestral lineage are discrete states in the continuous process, on the true tree these corresponds to actual cells but on the inferred tree they need not correspond to actual observed genotypes. Instead we can think about them as realizations along the continuous mutation process defined by the inferred tree. The COAR value is then a similarity measured between the true cell genotype and the inferred realizations, each sampled from the true and inferred mutation processes respectively, and in the case of a mismatch between the number of realizations and cells, a gap will be introduced in the alignment to compensate.

Using the aligned ancestral lineages it is now possible to derive a score, similar to a sequence alignment score. We use negative penalties for mismatches and zero points for matches, and furthermore normalize the alignment score to the smallest possible score (all mismatches) for that lineage, yielding the COAR value for a single lineage *i*:

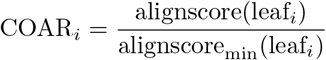

Where alignscore is the score of the alignment between the true and inferred ancestral lineages and alignscore*_min_* is the smallest possible score given the number and length of the sequences in the ancestral lineages. The alignment score is defined in terms of penalties, so all values are less than or equal to zero. Since both numerator and denominator are negative the COAR value is positive.

COAR is defined in the range from 0 to 1, where 0 is a perfect ancestral sequence reconstruction and 1 is the worst. The COAR value is comparable across different trees, methods and datasets because of this normalization. Its value can be interpreted as the average per-site error across all the inferred ancestral lineage sequences. COAR for a single ancestral lineage can be expanded to the tree level by calculating the mean COAR value for the whole tree:

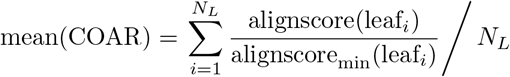

**Figure S21:**
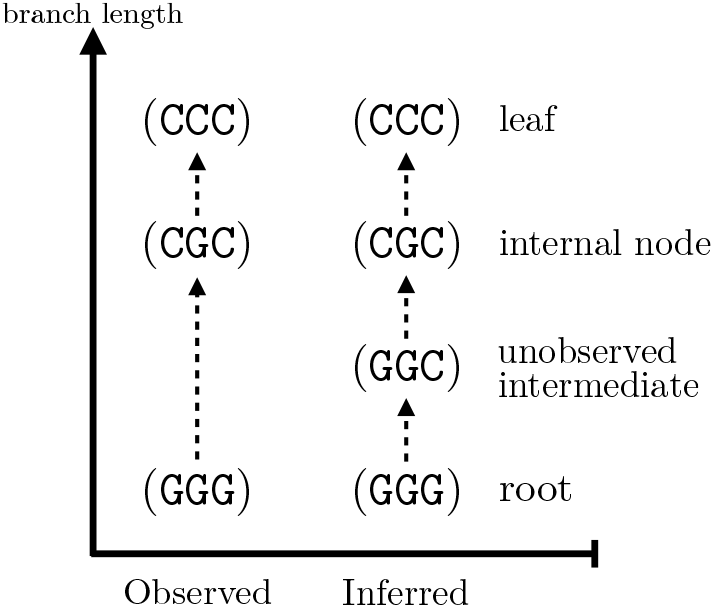
One interpretation of the COAR value is that it is the distance between the true and inferred mutation histories, here shown by the true and inferred ancestral lineage nodes of an example phylogeny. The true ancestral lineage (left side) represents actual observed cells where the genotype is a known constant. The inferred ancestral lineage (right side) represents the estimated genotypes at branching points along the inferred topology. In some cases there is a mis-correspondence between observed cells in the true phylogeny and the branching points in the inferred tree. These are treated as missing realizations and ignored in the alignment of the two mutation histories.

Where *N_L_* is the number of leaves on the tree.

#### Calculating COAR values - example with a known root

As an example of how the COAR metric works we will present a small example, summarized in Figure S20 with the light blue leaf chosen for lineage reconstruction and the true and inferred ancestral lineages marked in each tree with red dashed lines. The root sequence is a known state called the naive sequence. Assume that the true phylogeny is known with corresponding ancestral sequences. Now take a leaf sequence on the tree and reconstruct its ancestral lineage by extracting the parent, the parent’s parent, etc. until the root is reached, tabulated in Table S3. This ordered list of sequences constitute the reconstructed ancestral lineage for the chosen leaf and it always starts at the root and ends at the leaf, therefore we are imposing this as a restriction on the alignment. Furthermore, these two known states they do not count towards the COAR value.

**Table S3:** Reconstructed ancestral lineage for true and inferred trees as shown and marked by red dashed line in Figure S20.

|  | True | Inferred |
| --- | --- | --- |
| Naive (N) | AAA | AAA |
| $A_1$ | AAT | TAT |
| $A_2$ | ATT | - |
| Leaf (L) | TTT | TTT |

In the case of a wrongly inferred topology the true and inferred list of ancestral lineage sequences can have different length. It is therefore necessary to find a way of getting the best possible alignment between these two lists. We know the start and end of this alignment but the sequences in between are free to be shifted up or down to maximize the alignment fit. We adapt the Needleman and Wunsch dynamic program solution (82) to solve this as an alignment problem. A notable difference to the original algorithm is that it was intended to align two sequences of characters, like DNA or amino acids, while in this application a list of whole sequences are aligned.

The first step in the alignment algorithm is to calculate a score matrix of all pairwise sequence comparisons. For this example we use the negative Hamming distance as a score, however, the score function can be extended to reflect different situations, like imposing a larger penalty for non-synonymous rather than synonymous mutations. The score matrix is tabulated in Table S4.

**Table S4:** Score matrix based on all pairwise distances between the sequence in Figure S20.

| | N | $A_{1t}$ | $A_{2t}$ | L |
| --- | --- | --- | --- | --- |
| N | 0 | -1 | -2 | -3 |
| $A_{1i}$ | -2 | -1 | -2 | -1 |
| L | -3 | -2 | -1 | 0 |

Now we are ready to initializing the alignment grid used in the dynamic programming solution of the alignment problem. Initialization is started by inserting the scores of pure gap characters i.e. first row and first column (Table S5), and we enforce alignment of the two root sequences by setting these gap penalties to negative infinity. Similarly, we disallow introduction of gaps in the longest of the two lists, also by penalizing with negative infinity (Table S6). Setting certain gap penalties to negative infinity is a simple way of dealing with disallowed gaps and it also works well for implementations.

**Table S5:**
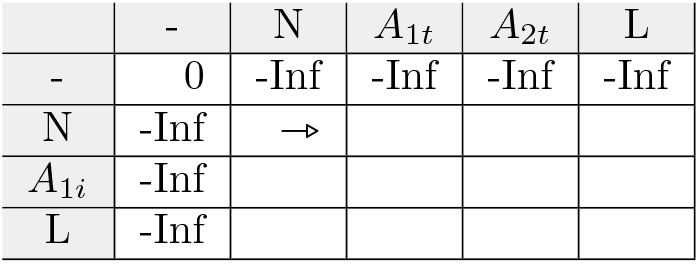
The starting alignment grid, initialized with negative infinite gap penalties to disallow gap opening in the beginning of the alignment. The grid is filled up from left to right row by row, starting in the cell marked by 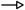.

| | - | N | $A_{1t}$ | $A_{2t}$ | L |
| --- | --- | --- | --- | --- | --- |
| - | 0 | -Inf | -Inf | -Inf | -Inf |
| N | -Inf | → | | | |
| $A_{1i}$ | -Inf | | | | |
| L | -Inf |  |  |  |  |

Then the alignment grid is filled up, starting with the cell marked by 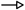 in Table S5, progressing to the rightmost cell and continuing in the same fashion on the next row. Cells are filled up using the following maximization:

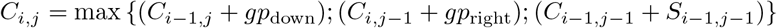

Where *C_i,j_* is the *i*th row and *j*th column cell in the grid, *gp*_down_ is the penalty of making a downwards gap, *gp*_right_ is the penalty of making a rightwards gap and *S_i_1,j_1_* is the score of aligning the *i*th, *j*th elements found by look-up in the score matrix (Table S4) In this example the longest list is that of the true ancestral lineage so in this list gaps are disallowed. In the inferred lineage gaps are allowed but not penalized: *gp*_down_ = –Inf and *gp*_right_ = 0.

The grid is filled and the final alignment score is the number in the rightmost bottom cell (Table S6).

**Table S6:** The filled alignment grid, ready for tracing back the best alignment. The rightmost bottom cell contains the score for the best alignment.

| | - | N | $A_{1t}$ | $A_{2t}$ | L |
| --- | --- | --- | --- | --- | --- |
| - | 0 | -Inf | -Inf | -Inf | -Inf |
| N | -Inf | 0 | 0 | 0 | 0 |
| $A_{1i}$ | -Inf | -Inf | -1 | -1 | -1 |
| L | -Inf | -Inf | -Inf | -2 | -1 |

The last step is to traceback the best path through the alignment grid and return this as the list alignment. The traceback starts from the leaf sequence, in the right bottom corner, and ends with the naive sequence in the left top corner. A diagonal step is equivalent to a sequence match, a left move is introducing a gap character in the inferred list and a move up is introducing a gap in the true list. The best path is found by progressively moving upwards, choosing the move with:

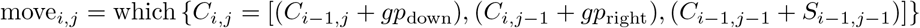

Notice that this path has already been generated when the alignment grid was filled up and could be cached. The resulting alignment and the penalty for each position is tabulated in Table S7.

Lastly the alignment score is normalized by the smallest possible alignment score i.e. no similarity between sequences in the lists. This normalized number is the COAR value and it runs between 0 to 1. In the presented example we only calculated the COAR value for the reconstructed ancestral lineage from one leaf, but by doing the calculations on all leaves on the tree and taking the average, the mean COAR value for the whole tree would be computed.

**Table S7:** The resulting alignment and the penalties for each position.

| True | N | $A_{1t}$ | $A_{2t}$ | L |
| --- | --- | --- | --- | --- |
| Inferred | N | $A_{1i}$ | - | L |
| Penalty | 0 | -1 | 0 | 0 |
| Max penalty | 0 | -3 | 0 | 0 |
| COAR | $-1/-3=0.333$ | | | |

